# Temporal Associations Between Human Milk Metabolites and the Infant Gut Microbiome and metabolome

**DOI:** 10.1101/2025.10.28.681130

**Authors:** Heidi Isokääntä, Laura Perasto, Santosh Lamichhane, Minka Ovaska, Teemu Kallonen, Eveliina Munukka, Matilda Kråkström, Hasse Karlsson, Linnea Karlsson, Henna-Maria Kailanto, Katri Kantojärvi, Tiina Paunio, Matej Orešič, Ulrik Kræmer Sundekilde, Alex M. Dickens, Anna-Katariina Aatsinki

## Abstract

**Introduction:** Human milk is vital in establishing a healthy infant gut microbiome. Gut microbes and their metabolites are important for host development. However, our understanding of how milk metabolites are connected to the gut microbiome and metabolome in early-life remains limited. We aimed to investigate associations between the human milk metabolome and the gut microbiome in infancy.

**Methods:** The fecal and milk samples were collected at 2.5 (n=283), 6 (n=129) and 14 (n=65) months of age. Gut microbiota was analyzed with amplicon sequencing, the gut metabolome was analyzed with gas chromatography-mass spectrometry and liquid chromatography-mass spectrometry, and milk metabolites were analyzed with proton nuclear magnetic resonance.

**Results:** Bifidobacterium and other butyrate producers were associated with ethanolamine and methionine level in the milk, in a time-dependent manner. Elevated milk fucosylated oligosaccharide, LNFP-I and caprylate concentrations were positively associated with secondary bile acid levels following the introduction of solid foods. Additionally, acidic oligosaccharides 3-SL and 6-SL were positively associated with long-chain carbohydrates prior to solid food introduction, however negatively associated thereafter.

**Conclusion:** Our results highlight that milk metabolites are associated with biologically relevant gut microbial metabolites. Only limited associations were observed between milk composition and the microbiome, which differed between early and late infancy. This suggests milk metabolome may differentially influence microbial metabolism depending on dietary diversity and/or the maturity of the microbiome.

**Importance:** It’s essential to raise awareness about breastfeeding and its variety of health benefits. Expanding our understanding of how human milk small molecules with potential bioactivity interact with the gut microbiome can help highlight these positive effects. Exploratory studies of human milk composition with gut microbial metabolites may support achieving this goal. Furthermore, understanding of biological effects of different components in human milk is important, and it can help to develop strategies to ensure optimal growth and development of an infant regardless of the breastfeeding status of the child.

## 1 Introduction

The infant gut microbiome undergoes dynamic developmental stages and is shaped by various early-life factors, including gestational age, delivery mode, and feeding practices [1–3]. Among these, human milk (HM) plays a central role in fostering a gut environment dominated by *Bifidobacterium* species. HM is rich in bioactive components—such as energy-yielding small molecules, enzymes, immunoglobulins, vitamins, and human milk oligosaccharides (HMOs)—that support microbial colonization and infant health [4, 5]. HMO composition is partly dictated by the maternal secretor genotype (FUT2), which is determined by fucosyltransferase-2 (FUT2) gene activity. Individuals harboring a non-functional FUT2 gene are termed non-secretors and they have markedly reduced levels of α1-2-fucosylated HMOs (e.g., 2ʹ-FL) compared to secretor mothers with functional FUT2 gene [6, 7]. HMOs are indigestible by the infant but serve as essential substrates for specific gut bacteria. These bacteria, in turn, generate microbial metabolites that act as signalling molecules, influencing immune function, neurodevelopment, and metabolic processes. [8–11] Another cohort study found associations between breastfeeding and bacterial genera in a longitudinal manner; breastfeeding supported *Bifidobacterium* and *Bacteroides* as early keystone organisms, directing microbiota development and consistently predicting positive health outcomes [12, 13]. In addition, they saw specific patterns in the associations between HMOs and infant gut genus abundances depending on the delivery mode, however the temporal changes in HMO profiles were unobserved [14].

Although it is relatively well-established how breastfeeding and HMOs are related to gut microbes, less is known about the metabolites produced by gut microbes. Microbial metabolites produced by gut microbes are often key mediators of health effects [15–17]. We and others have examined how feeding mode affects the composition of infant fecal metabolites, focusing on short-chain fatty acids (SCFAs) and secondary bile acids [18, 19]. Although, the shifts in the early-life fecal metabolome are driven largely by dietary alterations up to the first 2 years of life [20], there is still a lack of longitudinal research that simultaneously considers human milk composition, gut microbiota and fecal metabolite profiles. Understanding how natural variation in human milk influences microbiota assembly and function could guide strategies to promote healthy microbial development, support metabolic health, and mitigate disruptions such as those caused by antimicrobial exposures. To address this gap, we conducted a longitudinal cohort study involving infants to investigate the interconnected development of human milk metabolites, gut microbiota, and fecal metabolome across infancy.

In this study, we aimed to elucidate how HM metabolites, known to shape the infant microbiota, relate to fecal microbiota and metabolome at 2.5, 6 and 14 months. We hypothesized that HM metabolite composition and maternal/infant FUT2 status predict gut microbial composition among breastfed infants and that the differences are reflected in the fecal metabolite profile. We aim to study roles of that both maternal and infant FUT2 secretor status on infant gut microbiota and metabolome, since maternal FUT2 status (secretor status) alters in the HMO composition [21] and the infant FUT2 status alters the fucosylated mucin production [22–24], and thus these factors may have an effect on the microbiome, respectively. We expect to observe links between milk HMOs, increased level of bifidobacteria, short-chain fatty acids, and aromatic lactic acids [19], as well as lipid-class metabolites and fecal bile acids. Moreover, we hypothesize that associations between milk, fecal microbiota and metabolome may change during the first year of life. Results may reveal the variation in healthy microbiome development. This may be important for the foundation of gut microbiome and how the seeding effect of microbiome for later life health is formed.

## 2 Materials and Methods

### 2.1 Study participants and sample collection

Data was collected in FinnBrain birth cohort study [25] from the region of southwestern Finland and Åland in 2013-2016. Infant fecal samples (total 1002) were collected at 2.5 (n=444), 6 (n=256) and 14 (n=302) months of age in a preservative-free tube at home. They were delivered to a study visit and laboratory at +4C. The fecal samples were frozen at -80°C within 48h after sample collection. For fecal metabolomics, only samples that were frozen within 24h were included.

Human milk (HM) samples were collected at 2.5 (n=406), 6 (n=176) and 14 (n=80) months of age during study visits in the presence of a study nurse. Mothers were instructed to feed their infant from their right breast 1.5–2 h prior to the study visit but breastfeeding from the left breast was allowed based on infant needs during the same day. While wearing latex gloves, mothers were instructed to express manually 10 ml of foremilk into a sterile cup from their right breast. Samples with lower volumes were not excluded. After the sample collection, the sample was stirred and then divided into aliquots, stored at 4°C, and frozen at −80° during the same day.

Mothers filled questionnaires on background factors at 14 gestational weeks (GW), 24GW, 36 GW and on breastfeeding habits after delivery at 2.5, 6 and 14 months of infant age. Background questionnaires included information on maternal educational status and psychiatric symptoms with Edinburgh Postnatal Depression Scale (EPDS) [26] and Symptom Checklist 90 – anxiety subscale (SCL) [27]. Health records including information on parity, maternal pre-pregnancy body-mass index (BMI), birth date, delivery mode and perinatal antibiotic treatments were collected in from the hospital records of the Wellbeing Services of the County of Southwestern Finland.

### 2.2 Milk metabolomics

The milk samples were analyzed in Aarhus University, Department of Food Science. The samples intended for ^1^H nuclear magnetic resonance (NMR)-based metabolomics were processed in a random order as followed using standard protocol for milk-based metabolomics [28]. Samples were thawed in water bath and kept on ice while Amicon Ultra 0.5-ml 10-kDa spin filters (Millipore, Billerica, MA, United States) were being washed three times. The samples were skimmed by centrifugation at 4,000 g, at 4°C for 10 min, fat layer removed, and 500 μL of the skimmed milk transferred to individual Amicon Ultra 0.5-ml 10 kDa spin filters. Next, the skimmed milk was filtered by centrifugation at 10,000 g at 4°C, for 30 min and 400 μL of filtered milk from each sample was transferred to an individual 5-mm NMR tube. In each tube, 200 μL D2O with 0.05% 3-(trimethylsilyl)propanoic acid (TSP, Sigma-Aldrich, Saint-Louis, MO, USA) was added. Spectra acquisition was acquired according to the study of Sundekilde et al. [28]. Using a Bruker Avance Neo-IVD 600 spectrometer equipped with a 5-mm 1H BBI probe (Bruker BioSpin, Rheinstetten, Germany), ^1^H-NMR spectra were acquired at 298 K and a 1H frequency of 600.13 MHz. A standard metabolomics pulse experiment (Bruker pulse sequence: noesypr1d) was run to acquire one-dimensional spectra with a relaxation delay of 4 s. During the relaxation delay, water suppression was performed, and a total of 64 scans were comprised of 65.536 data points with a spectral width of 19.84 ppm. The resulting ^1^H-NMR spectra were all referenced to TSP signal at 0 ppm. A line-broadening function by 0.3 Hz was applied to each ^1^H NMR spectra, followed by a Fourier transformation. Preprocessing of ^1^H-NMR spectra was subsequently conducted by phase and baseline corrections, both automatically and manually using Topspin 3.2 (Bruker Biospin, Rheinstetten, Germany). Metabolite identification and concentrations was established using Chenomx NMR suite version 10.1 using the built-in and an in-house library of metabolite standards (Chenomx Inc., Edmonton, Canada)

Milk metabolite data consists of 44 compounds. Milk metabolites were grouped into five groups based on their chemical classification: HMOs, Energy metabolism, Amino acids & Protect nutrients, Bacterial fermentation and Lipids & fatty acids (Table S1). Milk samples give a snapshot of compound concentrations and therefore normalization was needed. Normalization for milk metabolites was done by lactose content since the lactose is relatively stable in mature milk and is correlated with milk volume [29] and has previously been used for normalization [28]. Data was log-transformed.

#### 2.2.1 Pheno- and genotypes of secretor status (FUT2)

Motheŕs secretor status of 2’-Fucosyllactose (FUT2) was determined by the concentration of 2’-FL in milk. Infant secretor status (FUT2) was estimated based on genetic variation (single nucleotide polymorphism, SNP) identified from DNA collected from blood samples as in previous study [30]. Briefly, DNA from cord blood was extracted as instructed by standard procedures at the Finnish Institute for Health and Welfare (THL). Extracted DNAs were genotyped with Illumina Infinium PsychArray BeadChip (Illumina, San Diego, CA) comprising 603132 SNPs at Estonian Genome Centre (Tartu, Estonia), and quality control was carried out with PLINK (www.cog-genomics.org/plink/1.9/) [31]. Markers were deleted for missingness (> 5%) and Hardy–Weinberg equilibrium (P < 1 × 10–6). Subjects were checked for missing genotypes (> 5%), relatedness (identical by descent calculation, PI_HAT > 0.2) and population stratification (scaled multidimensionally). Genotyped data was pre-phased with Eagle 2.3.5 [32] and imputed with Beagle 4.1 [33] using the Finnish population-specific SISu v2 imputation reference panel. One SNP of FUT2 gene rs601338 (non-functional FUT2 gene) was investigated to determine the secretor status of infant. The SNP was coded as follows: minor/minor=2 (A/A), major/minor=1 (G/A) and major/major=0 (G/G), meaning that subjects with a value of zero (homozygous for the major allele G) have a normally functioning FUT2 gene on both alleles (i.e., secretors), while all other genotype combinations were considered as non-secretors.

### 2.3 Gut microbiome

Only samples that were frozen within 48 h of sample collection were sequenced. For DNA extraction, 1 ml of lysis buffer was added, and the samples (∼100mg) were homogenized with glass beads 1000 rpm / 3 min. The samples were centrifuged at high speed (> 13000 rpm) for 5 min. The lysate (800μL) was then transferred to clean tubes. DNA was extracted using a semiautomatic extraction instrument Genoxtract with DNA stool kit (HAIN life science, Germany) and the extraction proceeded according to the manufacturer’s protocol. For all procedures related to microbiota analysis, DNA- and RNAse-free plastics were employed to prevent nucleic acid degradation.

DNA yields were measured with Qubit fluorometer using Qubit dsDNA High Sensitivity Assay kit (Thermo Fisher Scientific, USA). The DNA extraction and sequencing were performed in the University of Turku. Bacterial community composition was determined by sequencing the V4 region of 16S rRNA gene using Illumina MiSeq platform (Illumina, USA). The sequence library was constructed with an in-house developed protocol where amplicon PCR and index PCR were combined [34]. Positive control (DNA 7-mock standard) and negative control (PCR grade water) were included in library preparation and sequencing runs. DADA2-pipeline (version 1.14) was used to preprocess the 16S rRNA gene sequencing data to infer exact amplicon sequence variants (ASVs) [35]. The reads were truncated to length 225 and reads with more than two expected errors were discarded (maxEE = 2). SILVA taxonomy database (version 138) [36, 37] and RDP Naive Bayesian Classifier algorithm [38] were used for the taxonomic assignments of the ASVs.

### 2.4 Fecal metabolomics

Fecal metabolites were already measured as described in the previous study [39]. The order of the samples was randomized before sample preparation. An aliquot was freeze-dried prior to extraction to determine the dry weight. The second aliquot (∼50mg) was homogenized with beads and 20 μL of water for each mg of dry weight in the fecal sample.

Bile acids were extracted by adding 40 μL fecal homogenate to 400 μL crash solvent (methanol containing 62,5 parts per billion ppb each of the internal standards LCA-d4, TCA-d4, GUDCA-d4, GCA-d4, CA-d4, UDCA-d4, GCDCA-d4, CDCA-d4, DCA-d4 and GLCA-d4) and filtering them using a Supelco protein precipitation filter plate. The samples were dried under a gentle flow of nitrogen and resuspended using 20 μL resuspension solution (Methanol:water (40:60) with 5 ppb Perfluoro-n-[13C9] nonanoic acid as in injection standard). Quality control (QC) samples were prepared by combining an aliquot of every sample into a tube, vortexing it and preparing QC samples in the same way as the other samples. Blank samples were prepared by pipetting 400 μL crash solvent into a 96-well plate, then dried and resuspended as the other samples. Calibration curves were prepared by pipetting 40 μL of standard dilution into vials, adding 400 μL crash solution and drying and resuspending them in the same way as the other samples. The concentrations of the standard dilutions were between 0.0025 and 600 ppb. The LC separation was performed on a Sciex Exion AD 30 (AB Sciex Inc., Framingham, MA) LC system consisting of a binary pump, an autosampler set to 15 °C and a column oven set to 35 °C. A waters Aquity UPLC HSS T3 column with a precolumn with the same material was used. The flow rate was 0.5 mL/min, and the injection volume was 5 μL.

The mass spectrometer used for this method was a Sciex 5500 QTrap mass spectrometer operating in scheduled multiple reaction monitoring mode in negative mode. Data processing was performed on Sciex MultiQuant.

#### 2.4.1 Quantification of SCFA

We adapted and modified the targeted SCFA analysis from previous work [40]. Fecal samples were homogenized by adding water (10 μL per mg of dry weight as determined for the BA analysis) to wet feces, the samples were homogenized using a bead beater. Analysis of SCFA was performed on fecal homogenate (50 μL) crashed with 500 μL methanol containing internal standard (propionic acid-d6 and hexanoic acid-d3 at 10 parts per million (ppm)). Samples were vortexed for 1 min and filtrated using 96-Well protein precipitation filter plate (Sigma-Aldrich, 55263-U). Retention index (RI, 8 ppm C10-C30 alkanes and 4 ppm 4,4-Dibromooctafluorobiphenyl in hexane) was added to the samples. Gas chromatography (GC) separation was performed on an Agilent 5890B GC system equipped with a Phenomenex Zebron ZB-WAXplus column; a short blank pre-column of the same dimensions was also added. A sample volume of 1 μL was injected into a split/splitless inlet at 285°C using split mode at 2:1 split ratio using a PAL LSI 85 sampler. Mass spectrometry was performed on an Agilent 5977A MSD. Mass spectra were recorded in Selected Ion Monitoring (SIM) mode. The detector was switched off during the 1 min solvent delay time.

#### 2.4.2. Analysis of polar metabolites

Polar metabolites were extracted in methanol. The method was adapted from the method used by Lamichhane et al.[41]. Fecal homogenate (60 μL) was diluted with 600 μL methanol crash solvent containing internal standards (heptadecanoic acid (5 ppm) valine-d8 (1 ppm) and glutamic acid-d5 (1 ppm)). After precipitation the samples were filtered using the same filter plates as above. One aliquot (50 μL) was transferred to a shallow 96-well plate to create a QC sample. The rest of the sample volume was dried under a gentle stream of nitrogen and stored at -80 °C until analysis. After thawing the samples were again dried to remove any traces of water. Derivatization was carried out on a Gerstel MPS MultiPurpoe Sampler followed by injection. The automatic derivatization was carried out using the Gerstel maestro 1 software (version 1.4).

Gas chromatographic (GC) separation was carried out on an Agilent 7890B GC system equipped with an Agilent DB-5MS column. A sample volume of 1 μL was injected into a split/splitless inlet at 250°C using splitless mode. The mass spectrometry was carried out on a LECO Pegasus BT system (LECO). ChromaTOF software (version 5.51) was used for data aquisition. The samples were run in 9 batches, each consisting of 100 samples and a calibration curve. To monitor the run a blank, a QC and a standard sample with a known concentration run between every 10 samples. Between every batch the septum and liner on the GC were replaced, the precolumn was cut if necessary and the instrument was tuned.

The retention index was determined with ChromaTOF using the reference method function. The reference file contained the spectras and approximate retention times of the alkanes from C10 to C30 as determined manually. A reference method was implemented for every sample to determine the exact retention time of the alkanes. Untargeted data processing was carried out using MSDIAL (version 4.7). The identification was carried out using retention index with the help of the GCMS DB-Public-kovatsRI-VS3 library provided on the MSDIAL webpage. The results were exported as peak areas and further processed with excel where the results were normalized using heptadecanoic acid as internal standard and the features with a coefficient of variance of less than 30 % in QC samples were selected. Further filtering removed alkanes and duplicate features. The IDs of the features which passed the CV check were further checked using the Golm Metabolome Database.

### 2.5 Statistical analyses

The analyses were performed using R [42] environment, version 4.2.2, 2022. Libraries factoextra [43], mia [44], vegan [45], boot [46], MOFA2 [47–49] and ggplot2 [50] were used. P-values (two-tailed) smaller than 0.05 were interpreted as statistically significant. While multiple testing, adjusted p-values (false discovery rate FDR) smaller than 0.05 were interpreted as statistically significant.

**The covariates** were selected *a priori* based on expected causal relationships (Fig. S1). The covariates that were used in the models as categorical variables were breastfeeding (partial, full), parity (primipara, multipara), delivery mode (vaginal, section), and maternal education (low, mid, high). Continuous variables were maternal BMI, child’s birth month, gestational age and maternal distress. Maternal distress was calculated as the sum of scaled mean EPDS (14GW, 24GW, 34GW, 3 mo, 6 mo and 1y) and scaled mean SCL (14GW, 24GW, 34GW 3 mo and 6 mo). In cases of missing values, the mean sum score was calculated based on the available measurement points. Age terms in mixed model refer to the segments of the piecewise linear function used to model the (continuous) child’s age at the sampling time. The breakpoint, that is, where the line was allowed to turn (break), was at 6 months. This was selected since Finnish recommendations state that solid foods should be started at 4-6 months [51].

#### 2.5.1 Milk composition, community composition and fecal metabolome

Beta diversity calculations were based on the Bray-Curtis dissimilarity using relative abundances of the detected bacterial genera. Dissimilarity of milk metabolome was analyzed with Euclidean. PERMANOVA analysis was performed using the adonis2 function with 999 permutations. Both marginal and by terms effects were calculated. To observe the association of overall milk metabolome and to reduce dimensionality, principal components analysis (PCA) was conducted for milk metabolites. PCA was conducted using the function prcomp. The first three principal components (PC1-PC3) were used as independent variables in the PERMANOVA.

Dissimilarity of infant fecal metabolome was studied with Euclidean in a principal component analysis by infant FUT2 status and mother secretor status. Group difference was tested with the function adonis, an R implementation of PERMANOVA. Taxonomical diversity (Shannon index) was compared between secretors and non-secretors with two sample t-test in each timepoint separately.

#### 2.5.2 Differential abundance analysis

Differential abundance analysis (DAA) was conducted using the Wilcoxon signed-rank test between the genus level presence-absence data [52] and milk metabolites. The genera that had at least ten occurrences in both groups (presence/absence) were included in the analyses. P-values were adjusted for multiple testing using the FDR method. Analysis was stratified by timepoints (2.5mo, 6mo and 14mo). Additionally, DAA was conducted with ALDEx2 [53] which uses probabilistic modelling and considers the count-compositional nature of the data by including scale modelling by default both unadjusted and adjusted for covariates. The DAA was completed for each timepoint.

#### 2.5.3 Associations between milk and fecal metabolites

The associations between each milk metabolite and each fecal metabolite were examined using Spearman’s correlation. Fecal metabolites were log2 transformed with pseudocount of half minimum values. P-values were adjusted for multiple testing using the FDR method. The 95% bias-corrected and accelerated (BCa) bootstrap confidence intervals were calculated for the correlation coefficient (based on 1000 bootstrap samples). Analysis was stratified by timepoints (2.5mo, 6mo and 14mo).

Next the following linear mixed effects models were used to examine longitudinally the association between fecal metabolite and milk metabolites:

*M: fecal metabolites ∼ intercept + maternal BMI + birth month + parity+ gestational age + maternal distress + delivery mode + maternal education + age terms x milk metabolites,*

where fecal metabolites (polar metabolites, SCFAs, BAs) and milk metabolites were used in the model one at a time. Interaction between each milk metabolite and child’s age at the sampling time was included in the models. Covariates are described above in previous section.

Milk and fecal metabolites were scaled (mean=0, sd=1). P-values were adjusted for multiple testing using the FDR method.

#### 2.5.4 Multi-Omics factor analysis

Multi-Omics factor analysis (MOFA) was conducted using the MOFA2 function. MOFA discovers the principal sources of variation in multi-omics datasets [47–49]. Milk and fecal metabolites and genera level microbiome datasets were used in MOFA2 at once. Only the prevalent genera were used (detection=0.01, prevalence=0.1). Genera were examined using the centered log ratio (CLR) transformation with pseudo count one. Timepoints were treated as groups to enable comparing the sources of variability that drive each timepoint i.e., group. Fecal metabolites were log_2_-transformed.

## 3 Results

In the sample studied, 78% of infants were fully breastfed in the 2-month timepoint. In the 6-month timepoint, 71% were partially breastfed (Fig. 1A). In the 14-month-old, 14% were receiving breast milk. The overlap of milk and stool samples got lower by age due to decreasing breastfeeding (Fig. 1B). Approximately, 84% of mothers were secretors in the whole cohort, which is slightly higher than previously observed in other cohorts globally (75-80%) [54, 55]. Infant FUT2 genotype mostly matched mother’s secretor type (Fig. 1C). The background characteristics of study sample have been listed in each timepoint (Table S2.)

**Figure 1.**
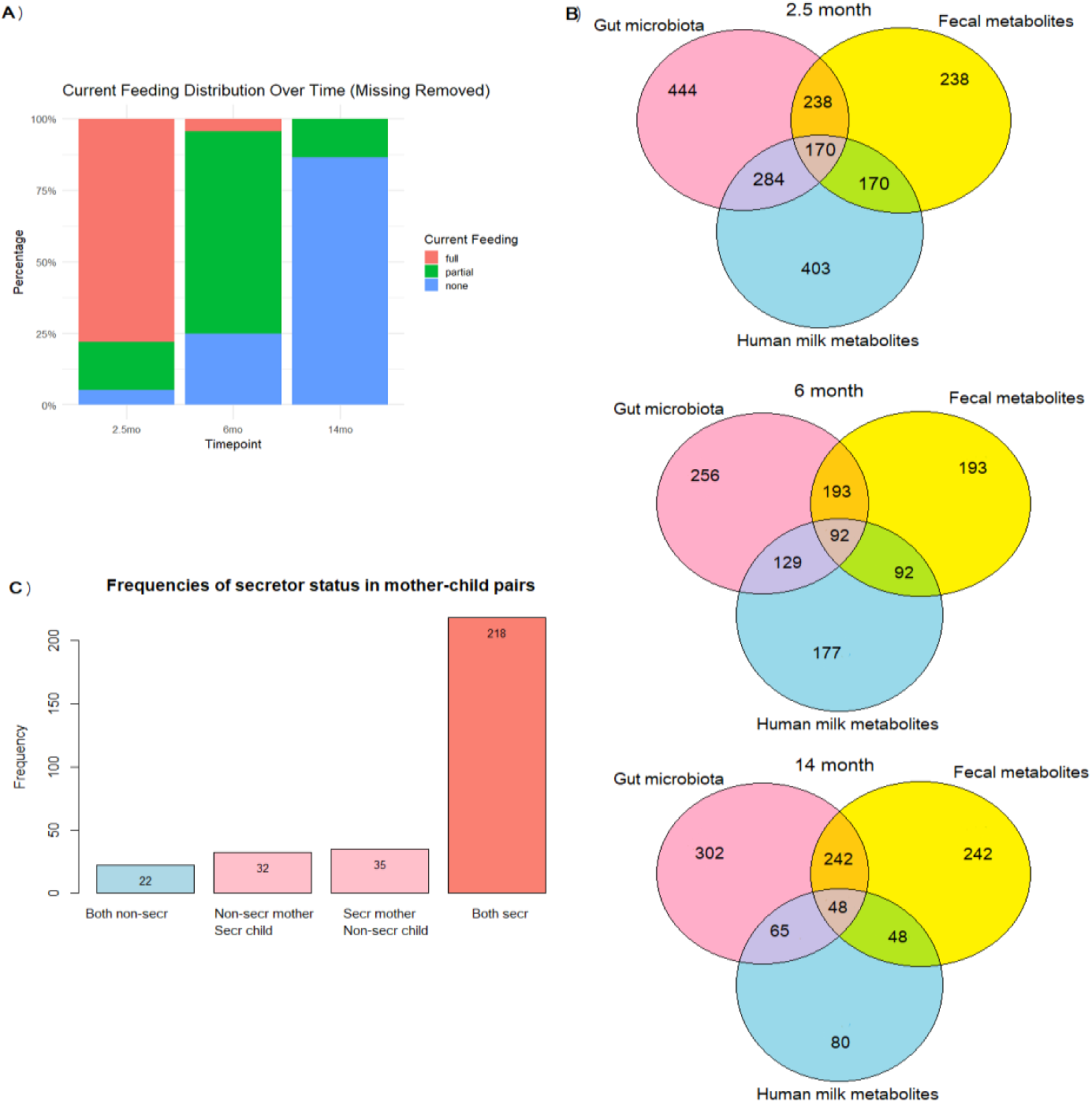
Overview of study samples. A) Breast feeding status in three timepoints. B) Overlap of stool and milk samples in multiomic setting across three timepoints. C) Frequencies of secretor status of mother-child dyads. 71% of dyads were both secretors.

### Diversity and dissimilarities in gut microbiome and milk metabolome

The gut microbiome got more uniform by time (Fig. 2A) and milk metabolome clustered according to mothers’ secretor status (Fig. 2B). Although the milk composition was clustered separately between secretors and non-secretors, the average amount of total HMOs was similar between the groups (Fig. S3). Total HMO concentration differed only in 2.5-month timepoint (mean concentration Secr. 16.9, Non-secr. 14.9, p<.001) by secretor status. HMO-profiles (Fig. S3) showed visually that non-secretors had more 3-FL while they were lacking fucosylated HMOs, i.e. 2’-FL (2ʹ-fucosyllactose), LNFP I (lacto-N-fucopentaose I), LDFT (lactodifucotetraose), and LNDFH I (lacto-N-difucohexaose I).

**Figure 2.**
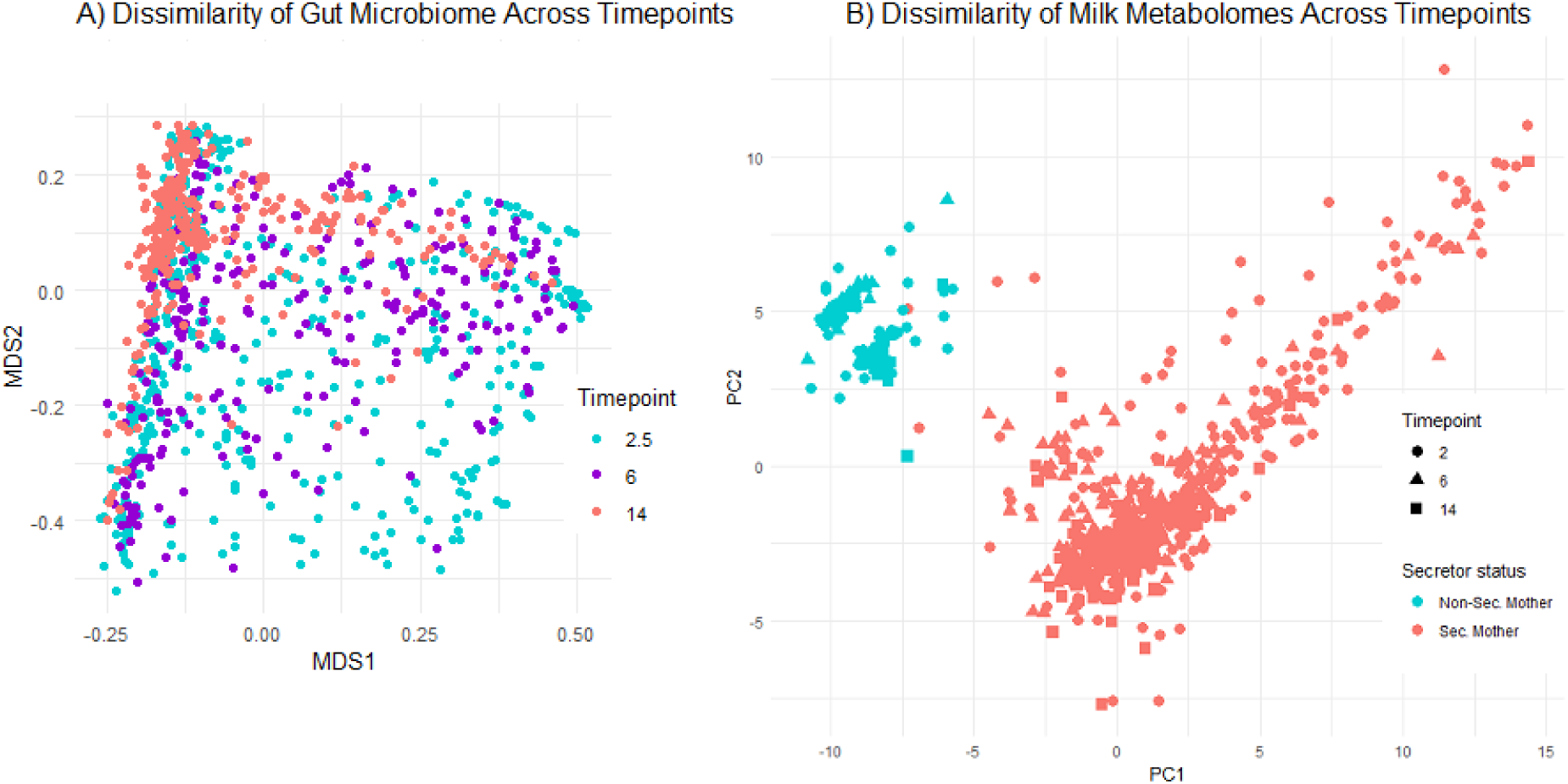
Dissimilarities. A) Infant gut microbiome beta diversity by Bray Curtis dissimilarity across three timepoints with multidimensional scaling. Microbiome became more uniform by time. B) Milk metabolomes by Euclidean and principal components by secretor status across timepoints. Timepoints are presented by shapes. Secretors are coloured with red and non-secretors with light blue.

The milk composition did not associate with the differences observed in the gut microbiome beta diversity in the infants, except PC1 of milk metabolome in 6 months explained a part of microbiome beta diversity (PERMANOVA, R^2^ = 0.02 p=0.02, Table S3). The taxonomic diversity (Shannon index) was not associated with either the secretor status of mother or the infant (t-test, p>0.4 in all timepoints) or the milk metabolome (Table S4). This indicates that the milk metabolome was not driving gut microbiota in the overall community composition or richness. Additionally, we examined whether fecal metabolome is driven by secretor status. Although fecal metabolome showed visually time-dependence (Fig. S4), no grouping of fecal metabolome by secretor status of mother or child were found (PERMANOVA p>0.05, Table S5).

### Milk metabolites and microbes in genus level correlate

In presence/absence analysis of microbes, there were many nominal significant differences in milk metabolite concentrations, but none withstood FDR correction (Table S6-S11). With ALDEx2, only two metabolite-microbe links remained after FDR correction: negative correlations between *Bifidobacterium* and ethanolamine/methionine (Fig. 3A). When considering nominal p-values, LNFP I was associated with *Bifidobacterium* in 6-month timepoint (Fig. 3B). Unidentified genus of *Enterobacteriaceae* associated positively with 3-SL in 2-month timepoint, while 3-FL associated negatively with Bacteroides. Overall, there were more associations in later timepoints showing more diverse microbiome composition. Glutamine had positive correlation with butyrate-producers *Faecalibacterium* and *Roseburia* [56] *in* 14-month timepoint (Fig. 3C), while 2’-FL correlated negatively with *Ruminococcus, Clostridioides and Citrobacter*. Sn-glycero-3-phosphocholine, a precursor in the synthesis of acetylcholine (a neurotransmitter), was associated in all three timepoints but with different genera. In the last timepoint, it was positively associated with *Bifidobacterium* and *Lactobacillus*.

**Figure 3.**
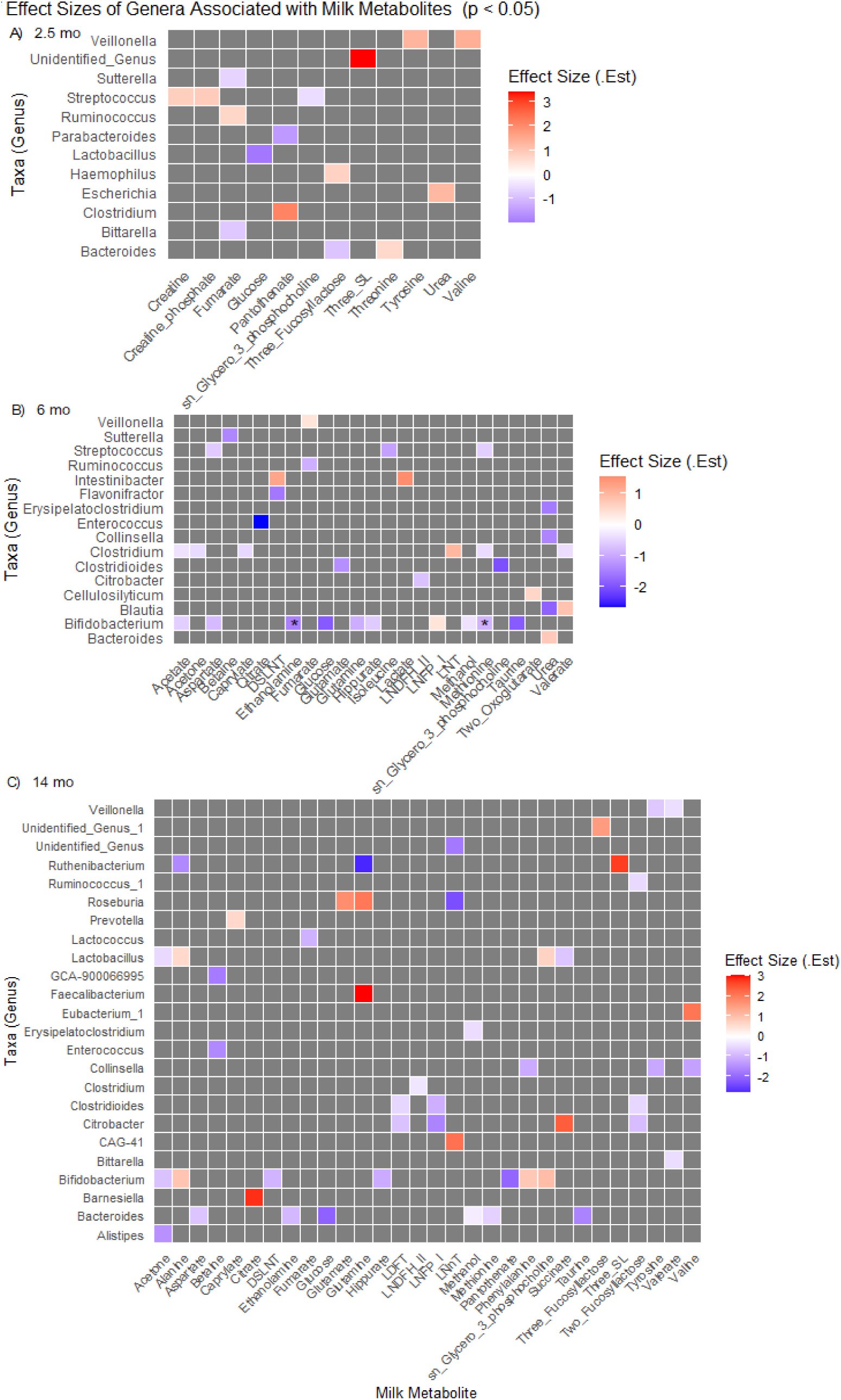
Differential abundances by ALDEx2 with milk metabolites across timepoints. All associations with unadjusted p-value <0.05 are presented here. Colours indicate direction and magnitude of estimates. Asterisks indicate adjusted p-values (FDR q-value) <0.05. Unidentified_Genus is an unknown genus of Enterobacteriaceae, Unidentified_Genus_1 is unknown Enterococcaceae, GCA-900-is unknown Oscillospirales and CAG-41 is from Clostridia class. A) estimates in 2.5 month timepoint. B) estimates in 6 month timepoint C) estimates in 14 month timepoint.

When adjusting for covariates using ALDEx2 with a linear model matrix, the associations between milk metabolites and microbial genera remained partially consistent (Fig. S5–S7). Notably, at 14 months, fewer genera showed significant correlations after adjustment, yet the effect estimates were stronger. For example, 2’-FL and 3-FL were negatively associated with *Ruminococcus*, while several milk energy metabolites showed positive associations with *Ruminococcus*. Interestingly, *Bifidobacterium* did not appear in the associations at either 6 or 14 months in the adjusted analysis.

### 3.3 Milk and fecal metabolites

Several associations (Fig. 4) were found between milk metabolites and fecal SCFA, however, the results did not withstand p-value correction. LDFT, which was missing with non-secretors, correlated positively with several fecal SCFAs, especially in 2.5- and 14-month timepoint. Milk amino acids had negative correlations to fecal acetic acid in 6-month timepoint. Milk amino acids associated with branched-chain fatty acids (Isovaleric and Isobutyric acid) in 2- and 6-month timepoints.

**Figure 4.**
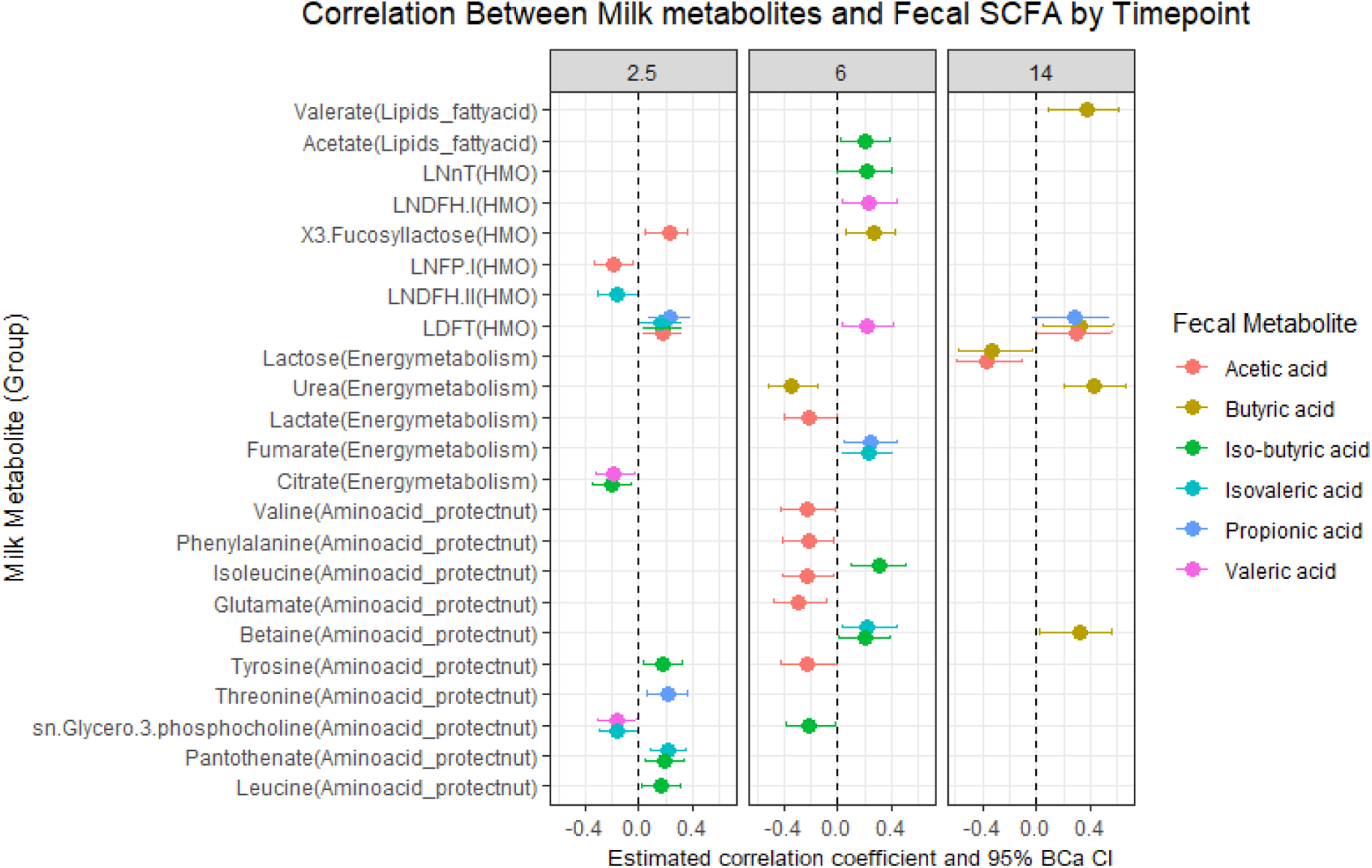
Milk metabolites correlated (Spearman) with fecal short chain fatty acids (SCFAs, see colors). These correlations have unadjusted p-values <0.05. The results did not withstand p-value correction (FDR). The 95% bias-corrected and accelerated (BCa) bootstrap confidence intervals were calculated for the correlation coefficient (based on 1000 bootstrap samples). Analysis was stratified by timepoints (2.5mo, 6mo and 14mo in gray panel).

Likewise, bile acids (BAs) were associated with several milk metabolites, but the results did not withstand p-value correction (Fig. 5). In 2.5-month timepoint, milk metabolites had positive associations with primary BAs, while 7-oxo-converted and secondary BAs were negatively associated. Interestingly, 2’-FL and 3-FL had opposite direction in BAs correlation.

**Figure 5.**
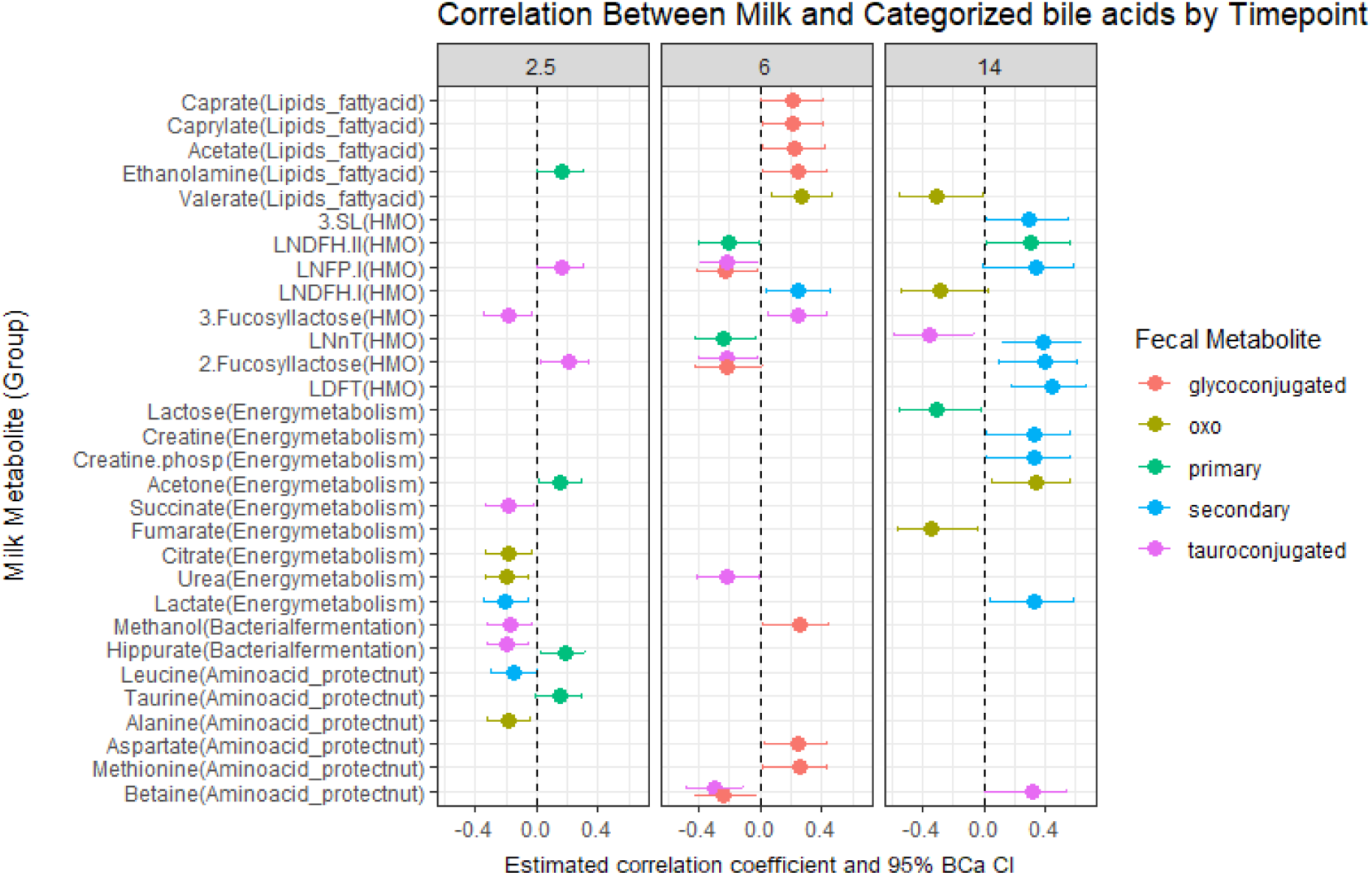
Milk metabolites correlated (Spearman) with fecal categorized bile acids (see colors). These correlations have unadjusted p-values <0.05. The results did not withstand p-value correction (FDR). The 95% bias-corrected and accelerated (BCa) bootstrap confidence intervals were calculated for the correlation coefficient (based on 1000 bootstrap samples). Analysis was stratified by timepoints (2.5mo, 6mo and 14mo in gray panel).

In 6-month timepoint, correlations were mainly in glycoconjugated BAs which were not seen in other timepoints. In 14-month timepoint, secondary BAs had several positive correlations. HMOs shifted the directions of correlation from 2.5month to 6month, i.e. 2’-FL and LNFP associated positively with tauroconjugated BAs in 2.5 months and negatively at 6 months, whereas an opposite pattern was observed for 3-FL.

There were high number of correlations (>700, p<0.05) between milk metabolites and fecal polar metabolites (Table S12). For instance, in 2.5 months milk caprate and caprylate, antimicrobial medium-chain fatty acids [57], correlated negatively with multiple fecal metabolites such as p-hydroxyphenyl lactic acid. Overall, associations were mainly negative and those were between milk fatty acids and fecal microbial and carbohydrate metabolism. However, in 2.5 months, when the most associations occurred, there were also positive associations between milk fatty acids (valerate, caprylate) and fecal polar metabolites, possibly originating from energy metabolism and xenobiotic sources.

### 3.5 Mixed model

In mixed model (adjusted for covariates mentioned in 2.5.3) we saw multiple associations between milk and gut metabolites when examining two time-intervals (2.5mo to 6 mo and 6 mo to 14mo) (Fig.6, Table S13). From fecal polar metabolites, long-chain carbohydrates fucose and ribonic acid had associations with milk 3-SL and 6-SL (Fig. 6C-D). Betaine and urea, which were already noted in DAA and Spearman, were associated positively with butyric acid in the later time interval (Fig. 6G-H). LDFT, which associated with multiple SCFAs with Spearman, correlated with 3,4-dihydroxyhydrocinnamic acid (likely xenobiotic) in both time intervals (Fig. 6E). LNFP I and caprylate correlated positively with secondary BAs in the later time-interval (Fig. 6A-B). Microbiota-derived 5-hydroxyindoleacetic acid, which has been seen to alleviate diarrhea [58], associated positively with milk sn-glycero-3-phosphocholine in the first time-interval. Nearly all correlations shifted direction after the first time-interval.

**Figure 6.**
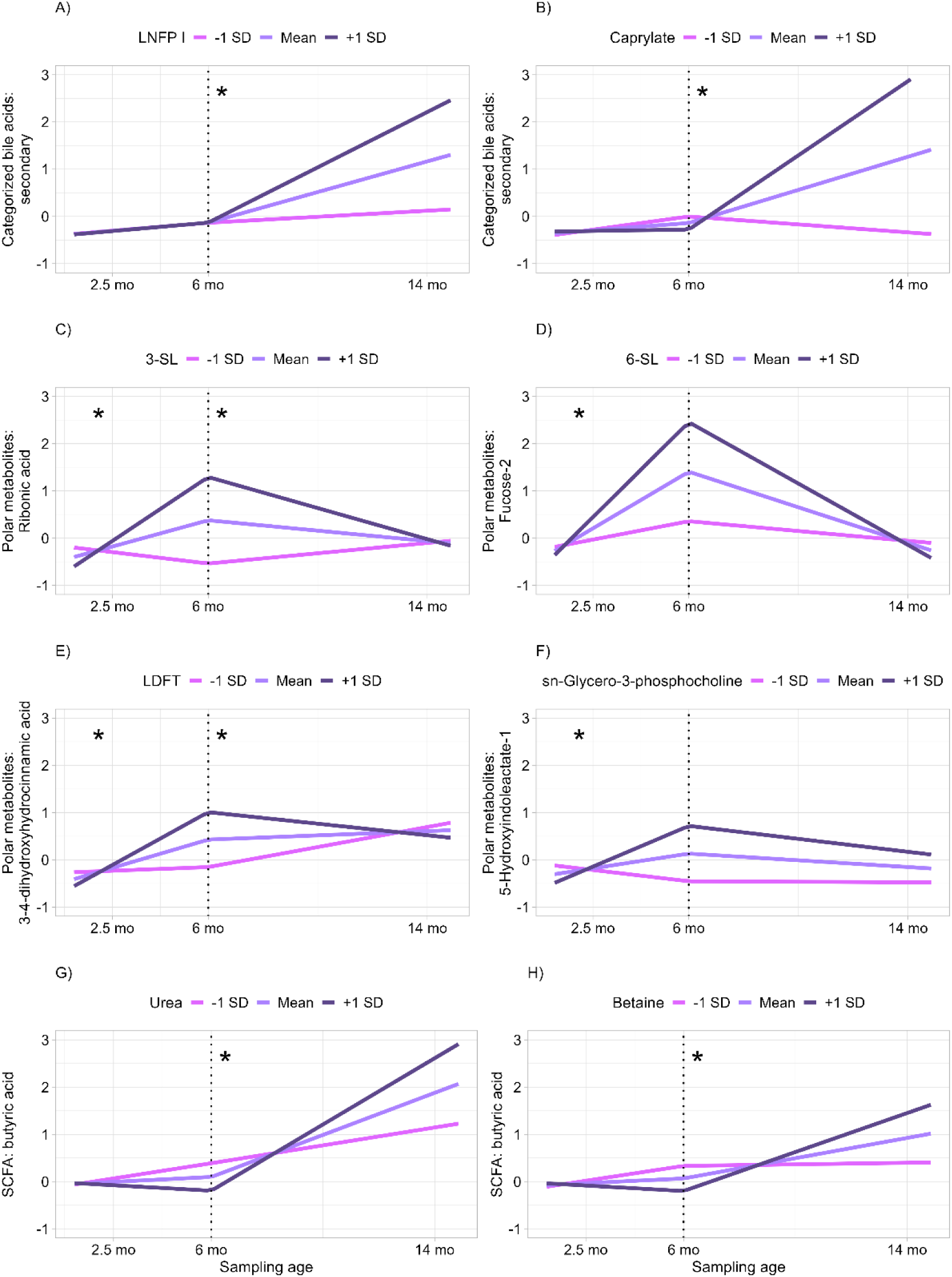
Mixed model by two time-intervals by sampling age. Associations between milk metabolite (on top of the plots) and fecal metabolites (y-axis, scaled transformed concentration) presented with mean values and standard deviations. Asterisks (*) indicate adjusted p-values <0.05.

### 3.6 Multi-Omics factor analysis

Multi-Omics factor analysis (MOFA, Fig. 7A.) with all omic-data sets was set out to see how much variance milk, fecal metabolites and gut bacterial genera explain in the data (Table S14) and to reduce features. Eight factors explained the variance and factors were loaded by different features (Factors 1-4 in Fig. 7B, all factors in Fig. S8). Overall, milk metabolites and stool-based omics explained variance in different latent factors, which capture underlying patterns of variation across omic data sets. With milk metabolites, variance was mainly explained by factors 1 (21.4%), 3 (23.3%) and 7 (10.6%) in 2-month timepoint and those remained largest throughout the timepoints. Those factors were loaded by milk energy metabolites, HMOs and fecal propionic acid. In bile acid data set, variance was explained by latent factor 2 (20.9%) and to some extent by 4 (5.2%) and 5 (7.0%) in 2-month timepoint and those factor values fluctuated by time. Other assays explained variance to lesser degree. Factor 1 was the only shared factor between milk and fecal metabolites, and it explained minor part of SCFAs (2.5mo: 1.5%; 6mo: 1.7%).

**Figure 7.**
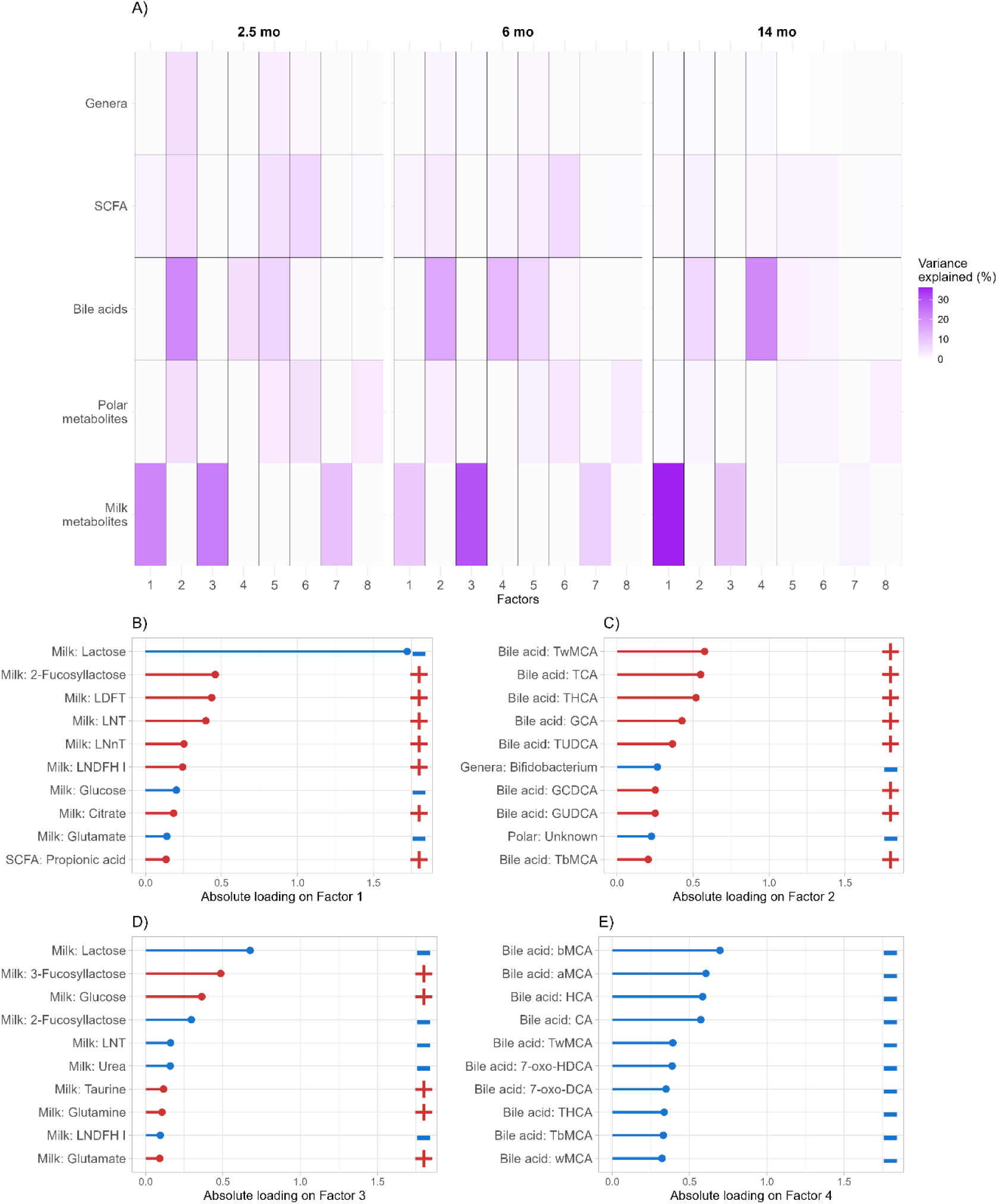
A) Multi-omic factor analysis (MOFA) with five data sets and 8 factors in three timepoints. B)-E) Loadings of factors 1, 2, 3 and 4. Top 4 factors and their top 10 features are presented here. Colors (red/blue) and +/- indicate the direction of the loadings.

Additionally, individual factor points were compared with breastfeeding (BF) variables (Fig. S9). Overall, categories of current breastfeeding were associated with factor 1 and factors 5 to 8 mainly at 2.5 months, while secretor status (mother) associated with factors 1, 3 and 5 (Fig. S9). Specifically, factors 5 and 6 were higher with full breastfed at 2.5mo timepoint (Fig. S9). Factors 1 and 5 were elevated in secretors at 2.5 and 6 months, respectively, while factor 3 was lower in 2.5 and 6 months (Fig. S9). Although both maternal secretor status and breastfeeding related to factor 5, these were at different timepoints.

Factors 1 and 3 were loaded mainly by HMOs and negatively by lactose (Fig. S8). Factor 1 was positively loaded with fucosylated HMOs, while factor 3 was positively loaded by 3-FL, milk glucose and milk amino acids and negatively by fucosylated HMOs (Fig S8). Factor 5 was negatively loaded by conjugated bile acid (taurine), fecal butyrate and positively with *Bifidobacterium*. Factor 6 was positively loaded with fecal sugars and negatively loaded with propionic acid (Fig S8).

## 4 Discussion

This study was set out to investigate the association between human milk metabolites and infant gut metabolites and microbiota. It is well known that human milk fosters infant healthy gut microbiome which produce beneficial microbial metabolites. However, there is a lack of longitudinal research that simultaneously considers human milk composition, gut microbiota and fecal metabolite profiles.

As expected, the milk HMO composition was influenced by mother’s secretor status, primarily due to the absence of specific fucosylated HMOs such as 2’-fucosyllactose (2’-FL), lacto-N-difucohexaose I (LNDFH I), and lacto-difucotetraose (LDFT). In addition, non-secretor mothers had higher concentration of 3-FL. In the mammary gland, FUT2 encodes an α1,2-fucosyltransferase for the synthesis of 2’-FL, while FUT3 gene is responsible for 3-FL production [21]. 3-FL has been also found to have prebiotic property, immunomodulatory effect, antiadhesive antimicrobials, antiviral ability, and gastrointestinal protection [59]. However, 2’-FL and 3-FL had opposing associations with fecal metabolites, such as secondary BAs. Although 2’-FL and 3-FL have been found to have similar properties, our results suggest that they also might differ in their relation to gut microbial metabolism. Hence, it can be speculated that the compensation of higher 3-FL may not address the deficiency of 2’-FL in the milk of non-secretor mothers in relation to gut metabolism. Moreover, one controlled trial showed that infants fed breastmilk or 2’-FL supplemented formula had similar systemic levels of microbial secondary BAs [60].

In our results, neither maternal or infant secretor status (FUT2) were associated with infant gut microbiota alpha and beta diversities. Although they have been considered important for gut health [55, 61, 62], our results align with recent studies indicating modest associations at best [63–65]. Moreover, in our results, the associations between the overall milk composition and infant gut microbiome diversity and community composition were mostly non-existent. Still, we observed an association between the first principal component of the milk and gut microbiota beta diversity at 6 months. Previous study showed that the proportion of human milk in the infant diet at 6 months was the prominent determinant of infant gut microbiota diversity [66]. One animal study reported that mother secretor status differentiates gut microbial beta diversity in one dimension, but HMO supplementation did not. [67].

We found that individual genera abundances were related to human milk composition. Fucosylated pentasaccharide LNFP-I was positively correlated with *Bifidobacterium* at 6 months and negatively with *Clostridioides* and *Citrobacter* at 14 months. LNFP-I is known to be consumed by *Bifidobacteria*, and can protect from enteropathogens [68] Additionally, in our study LNFP I exhibited a positive correlation with secondary BAs in later time interval, possibly indicating gut maturation [18]. Further, sialylated DSLNT in 2.5 months associated positively with *Veillonella* and negatively with *Escherichia* and further in 6 months with *Hungatella* and *Citrobacter*. *Escherichia* may also include possible pathogenic species. In a rat model of necrotizing enterocolitis, DSLNT was found to provide protection to necrotising enterocolitis [69], possibly by limiting the pathogenic bacteria. Interestingly, 2’-FL did not associate with *Bifidobacterium* but fucosylated HMOs LDFT, LNFP-I and 2’-FL associated negatively with *Clostridioides*, *Citrobacter* and *Ruminococcus* at 14 months. Hence HMOs appeared to limit potentially opportunistic pathogens [70–72]. Although the links between bifidobacteria and HMOs are well-documented [9, 73], previous study showed that also other bacteria have the capacity to metabolize HMOs and the by-products of HMO metabolism [9, 74]. This may underline our observation that HMOs were also related to other bacteria than *Bifidobacterium*, including members of Bacillota.

In addition to associations with bacterial genera, HMOs associated with bile acids, sugars and a xenobiotic metabolite (DHCA). 3-SL and 6-SL, both containing sialic acid bound with lactose, associated with higher fecal ribonic acid and fucose concentrations, respectively. The associations in 6-SL and fucose was apparent before weaning and 3-SL and ribonic acid associated also after weaning. Of note, 3-SL was also positively associated with unidentified genera in *Enterobactreriaceae* at 2.5 months. Previous research has demonstrated that sialylated HMO supplementation can increase transcription of genes related to monosaccharide and carbohydrate metabolism in *E. coli* and *Bacteroides fragilis* [75]. In this light, our finding can reflect that increased simple sialylated HMOs can result in higher abundance of members of Enterobacteriaceae and more extensive saccharolytic metabolism and higher concentration of carbohydrate derivates in stool.

Additionally, milk sn-glycero-3-phosphocholine (GPC), a breakdown product of phosphatidylcholine and a bioavailable choline source, was negatively associated with *Streptococcus* in 2-month-olds and *Clostridioides* in 6-month-olds and positively with *Bifidobacterium* and *Lactobacillus* at 14 months. GPC may shape the infant gut microbiota since some bacteria possess the necessary enzymes to break down GPC into its constituent parts, which can then be further metabolized for growth [76]. However, the effects on host health are uncertain since certain bacteria can utilize unabsorbed choline (e.g. some *Clostridium* and *Escherichia* strains) potentially to produce trimethylamine (TMA) which may further end up in host liver as TMAO, a potentially unfavourable substance for cardiometabolic health. [77, 78]

In our cohort the higher milk amino acid levels were linked to higher amount of branched short-chain acid (BCFA) isobutyric acid in 2.5-month-olds, but not in later timepoints. Previous studies have linked higher protein intake and amino acid metabolism to increased BCFA concentrations [79, 80]. Although amino acids are typically efficient at absorbing in small intestine, it may be that some escape to colon for bacterial metabolism if they are abundant in the diet. Interestingly, while isobutyric acid is metabolized from valine and isovaleric acid is produced from leucine [81], we observed positive correlations between isobutyric acid and leucine and tyrosine at 2.5 months as well as isoleucine at 6 months. In addition, milk amino acids, including valine, phenylalanine, isoleucine, tyrosine and glutamate, were negatively correlated fecal acetic acid at 6 months. BCFA are less studied in comparison to traditional SCFA, and whether our observation related to more complex bacterial cross-feeding in the intestines, remains an open question.

Urea and betaine were linked to higher butyric acid especially after weaning. Urea is a major source of nitrogen in the human milk, which may affect the gut homeostasis and bacterial metabolism [82, 83]. While the human host does not encode urease, which is an enzyme responsible for urea hydrolyzation. However, multiple microbes encode this enzyme, including *Bifidobacteria* and *Lacnospiraceae* [82]. Interestingly, the commensal bacteria with ureasis activity can use urea to produce SCFA, including butyrate [83]. Human milk betaine, primarily originating from dietary sources, has been associated with normal growth patterns in healthy infants as opposed to accelerated growth [84]. In addition, the betaine supplementation in early life in rodents and higher milk betaine in humans was related to increased *Akkermansia* abundance. Although *Akkermansia* had too low prevalence in our sample, it is known that *Akkermansia* is a SCFA producer [85]. Previous study has associated higher betaine levels in milk with lower adiposity and improved glucose homeostasis in adulthood, as observed in a mouse model [84]. These results highlight that also other human milk components than HMOs, such as urea and betaine, are associated with key microbial metabolites, including butyrate, and this link between human milk and gut microbiome is likely important for infant health [84].

We observed that milk lipids such as caprylate associates positively with glycoconjugated and secondary BAs, especially after weaning. Caprylate is a medium-chain fatty acid that typically increase during lactation [86] that is absorbed from the small intestines and serves as an energy source for the infant. Often lipid-containing diet increases the bile production [87, 88]. Host produces bile acids and conjugates them with taurine or glycine, and the conjugated BAs are deconjugated by bacteria and further metabolized to secondary BAs. The observed associations may reflect increased bile production subsequent microbial metabolism of BAs in response to higher lipid content in the milk. Bile acids are important physiological modulators that often undergo microbial metabolism [18], and our results suggest that milk lipids may be one factor affecting the bile acid enterohepatic circulation in infancy.

Our multi-omic analyses reflected that the data from different samples related to separate multi-omic factors, i.e. fecal data explained data-driven variance different from milk metabolome. This aligns with the observation that overall milk metabolome composition was not related to gut microbiota or metabolome overall composition. On the other hand, the factor loadings were related to breastfeeding and secretor status of the mother. Full-breastfed infants in 2.5 months, compared to partial feeding, had higher score in factors loaded positively by 7-oxo-HDCA, wMCA, and *Bifidobacterium*, which can be considered indications of healthy gut microbiome maturation [89–91]. On the other hand, there were negative loadings in butyric acid and other BAs, such as tauro- and glycine conjugates. This supports our earlier findings when we demonstrated that *Bifidobacterium* abundances were associated negatively with tauroconjugated BA concentration only in breastfed infants. Our results also corroborate earlier observation that 7-oxo-converted BAs correlated positively with *Bacteroides* and *Escherichia* in breastfed babies [39].

### Limitations and strengths

All milk samples were taken in a repeatably manner during study visits during daytime. Limitations of milk data concern the milk sampling (foremilk or hindmilk) and timing which may affect the milk composition. For that reason and the nature of lactation, collected milk samples represent snapshot of the milk. Moreover, metabolomics was performed using different analytical platforms — NMR for milk and GC- and LC-MS methods for fecal metabolites. While this approach may limit direct comparability of results, it also enables complementary coverage of distinct metabolite classes and enhances the overall breadth of metabolic profiling suitable for the sample type. The strength of this study is the unique longitudinal sample collection with multiomic approach. However, a part of data posed a challenge due to the high number of features which may explain high adjusted p-values. However, we observed similar relationships between different analysis methods, including cross-sectional correlations, DAA, multi-omic analyses and mixed models.

## 5 Conclusions

Our study uncovers dynamic, time-dependent relationships between human milk metabolites, the infant gut microbiome, and fecal metabolites. These associations varied between early and late infancy, suggesting that the influence of milk composition on microbial metabolism shifts with dietary diversification and microbiome maturation. While the overall milk composition was not related to gut microbiota or metabolome overall composition, our results highlight the relation between milk HMOs, amino acids, urea, lipids and key fecal metabolites such as butyrate and bile acids. Together, these insights advance our understanding of how human milk relates to the developing gut microbial metabolism and lay the groundwork for strategies to support optimal early-life health.

## Ethical evaluation

The study has a permit from the ethical committee of Hospital District of South-west Finland (ETMK: 57/180/2011). FinnBrain parents have signed a consent form about their children’s participation in research and given permission to use their samples for scientific purposes. Samples went through the laboratory process anonymously with research code to protect participants’ privacy.

## Conflict of interest/Disclosure statement

HMK is senior researcher at IFF. EM has previously worked at Biocodex. Other authors have no conflict of interest.

## Author Contributions

Conception and design of the work: HI, AKA, US, SL and AD. Acquisition, analysis, or interpretation of data: US, LK, HK, HMK, HI, TK, LP, MO, AD, AKA and SL. Drafting and substantial revision of the work: AKA, SL, HI, AD and LP. All authors have approved the submitted manuscript.

## Acknowledgements

We thank the FinnBrain Birth Cohort Study participants and personnel, Turku Metabolomics Center for the assistance and resources in the analyses of fecal metabolites. NMR data of milk was generated using research infrastructure at AU supported by FOODHAY (Food and Health Open Innovation Laboratory, Danish Roadmap for Research Infrastructure).

## Funding

We acknowledge several funders for making it possible to carry out this study. Finnbrain Birth cohort Study (HK) has been funded by Research Council of Finland (grant numbers 253270, 134950), Jane and Aatos Erkko Foundation, as well as Signe and Ane Gyllenberg Foundation. LK was funded by the Research Council of Finland (grant number 308176 and 325292), Yrjö Jahnsson Foundation (6847, 6976), Signe and Ane Gyllenberg Foundation, Finnish State Grants for Clinical Research (P3654), Jalmari and Rauha Ahokas Foundation, and Waterloo Foundation (2110-3601). HI was supported by the Finnish cultural foundation [no 00230482], Juho Vainio’s foundation and Doctoral school of clinical research of university of Turku. AKA and HI were supported by the Ane and Signe Gyllenberg Foundation. AKA was supported by Yrjö Jahnsson Foundation, Psychiatry Research Foundation, Emil Aaltonen Foundation, Brain Foundation, Instrumentarium Science Foundation, Duodecim Finnish Medical Society, Juho Vainio Foundation and the Research Council of Finland (grant number 347640). HMK was supported by the Finnish Cultural Foundation. SL was supported by the Research Council of Finland (decision number 363417). AMD has been funded by the Waterloo foundation and Research Council of Finland (347924). Further support was received by the “Inflammation in human early life: targeting impacts on life-course health” (INITIALISE) consortium funded by the Horizon Europe Program of the European Union under Grant Agreement 101094099 (to MO) and InFLAMES Flagship Programme of the Research Council of Finland (decision number: 337530). HMK was supported by the Finnish Cultural Foundation.

## Data availability statement and deposition

Due to Finnish national legislation and study participant rights, the individual-level data cannot be made available online, but data can potentially be shared with Research Agreement as part of research collaboration. Requests for collaboration can be sent to the Board of the FinnBrain Birth Cohort Study; please contact Linnea Karlsson.

## Supplementary figures

**Figure S1.**
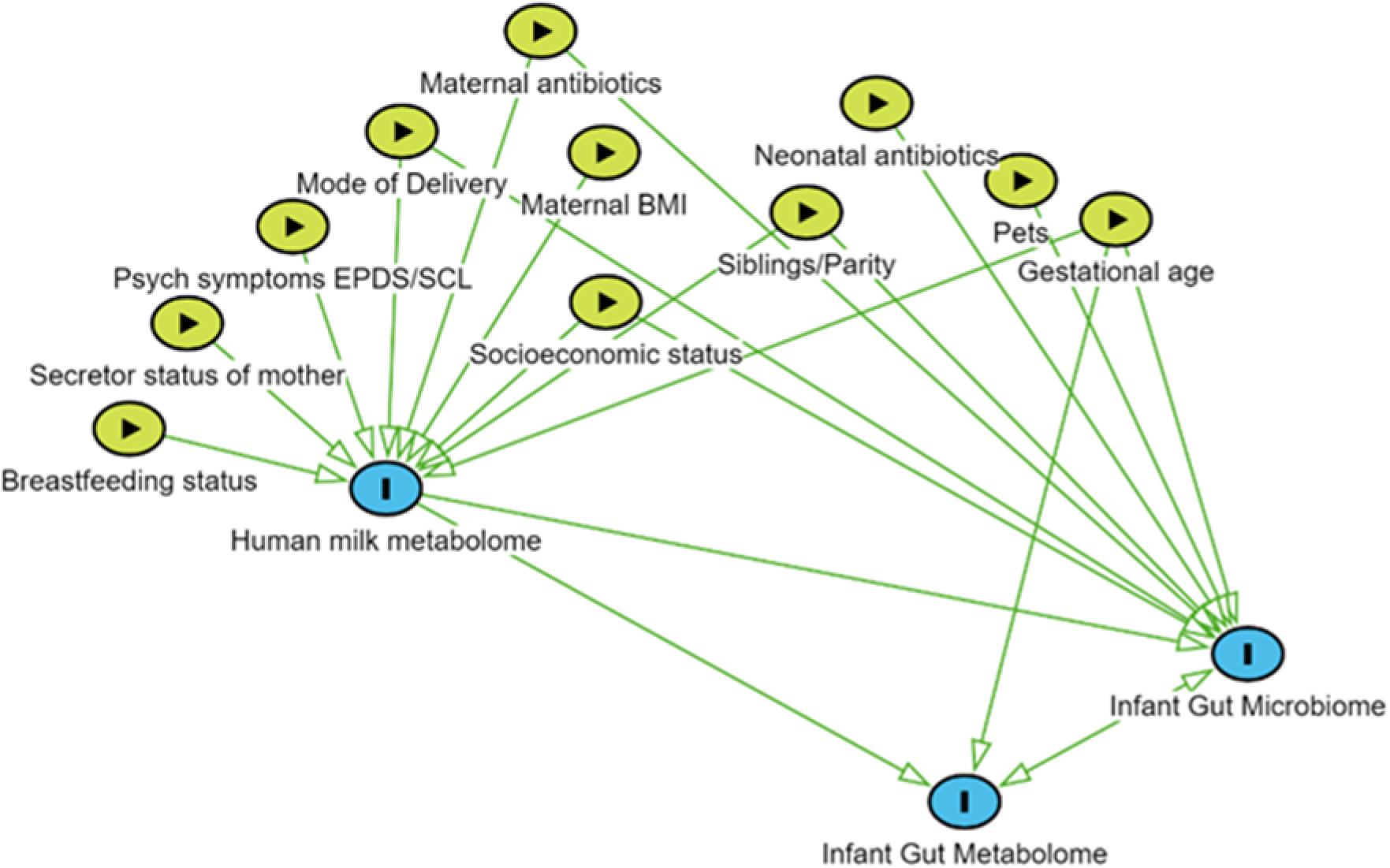
Exposures known to associate with human milk, gut microbiome and gut metabolome.

**Figure S2.**
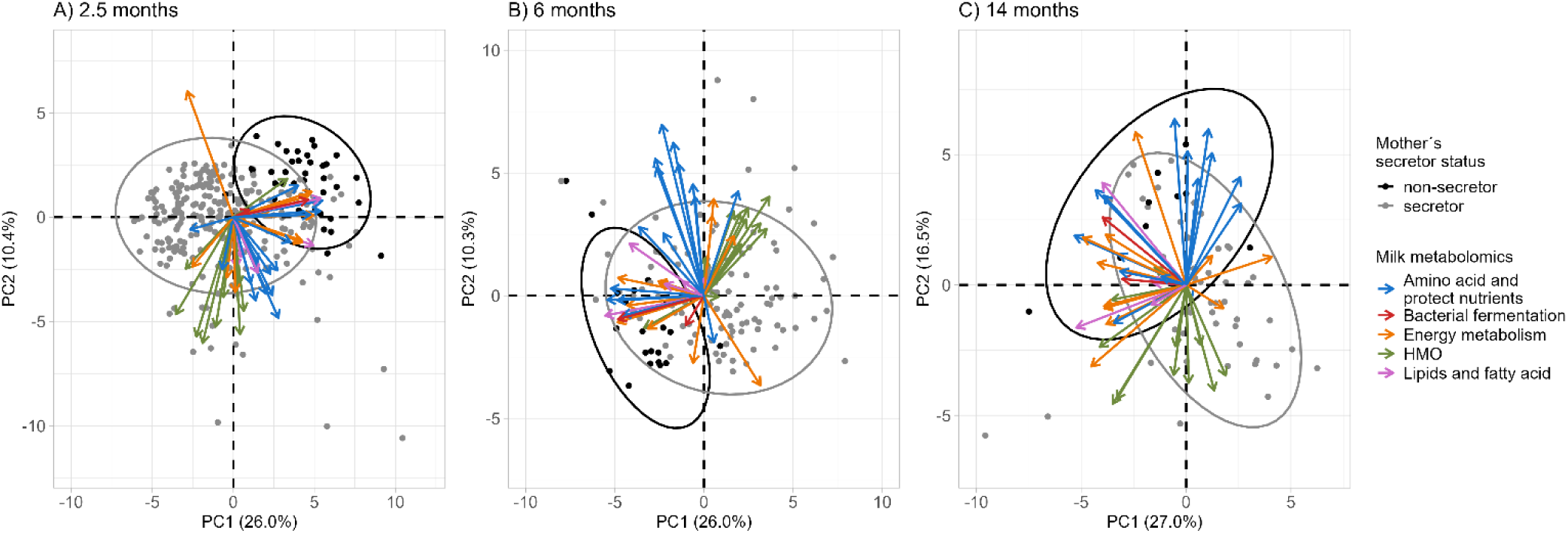
Milk metabolome dimensions and directions by metabolites in non-secretors (grey) and secretors (black) in three timepoints. Secretor status is sourced from 2’-FL concentrations in mother’s milk.

**Figure S3.**
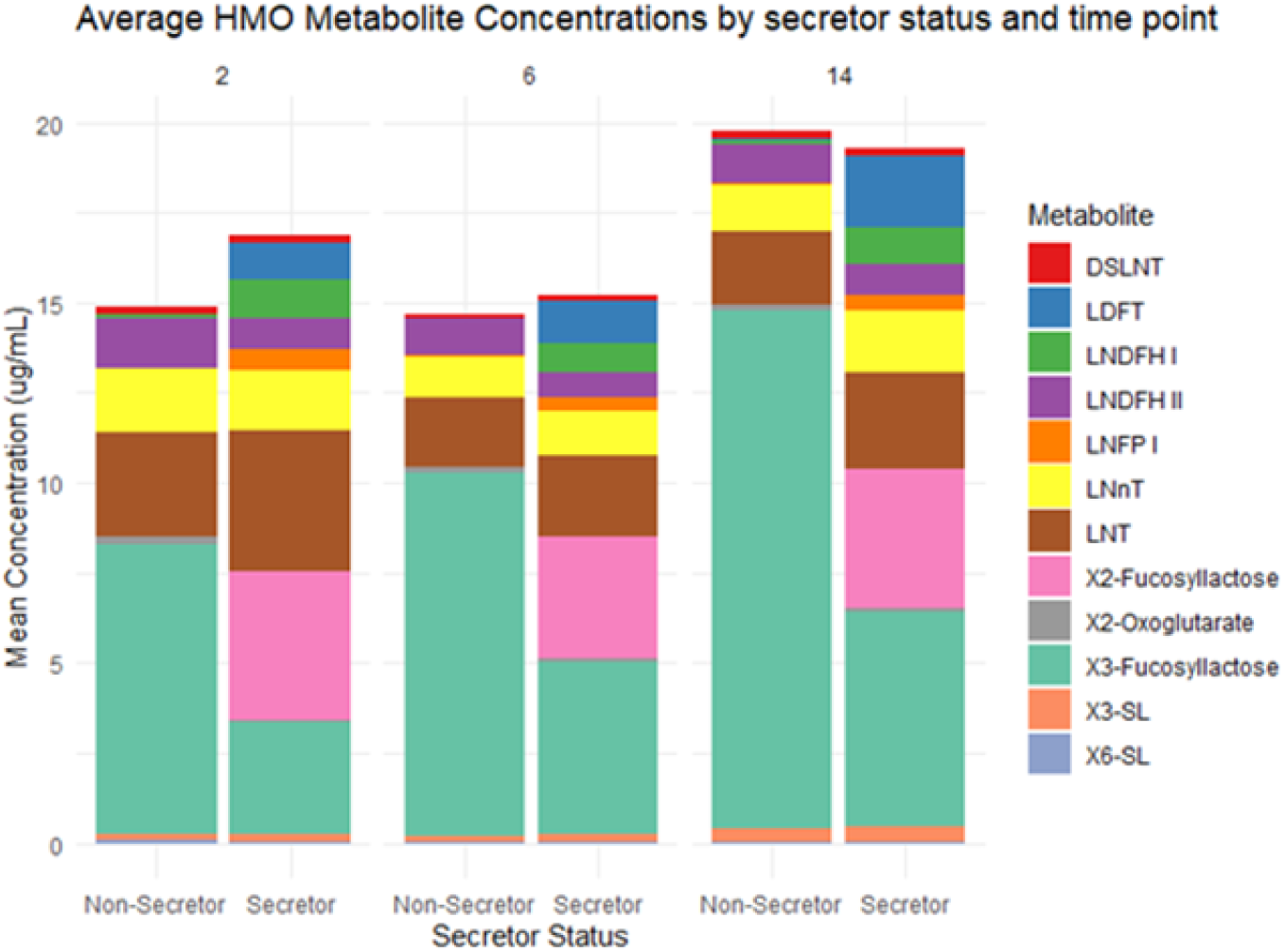
Mean concentrations of HMOs by mother secretor status and timepoint.

**Figure S4.**
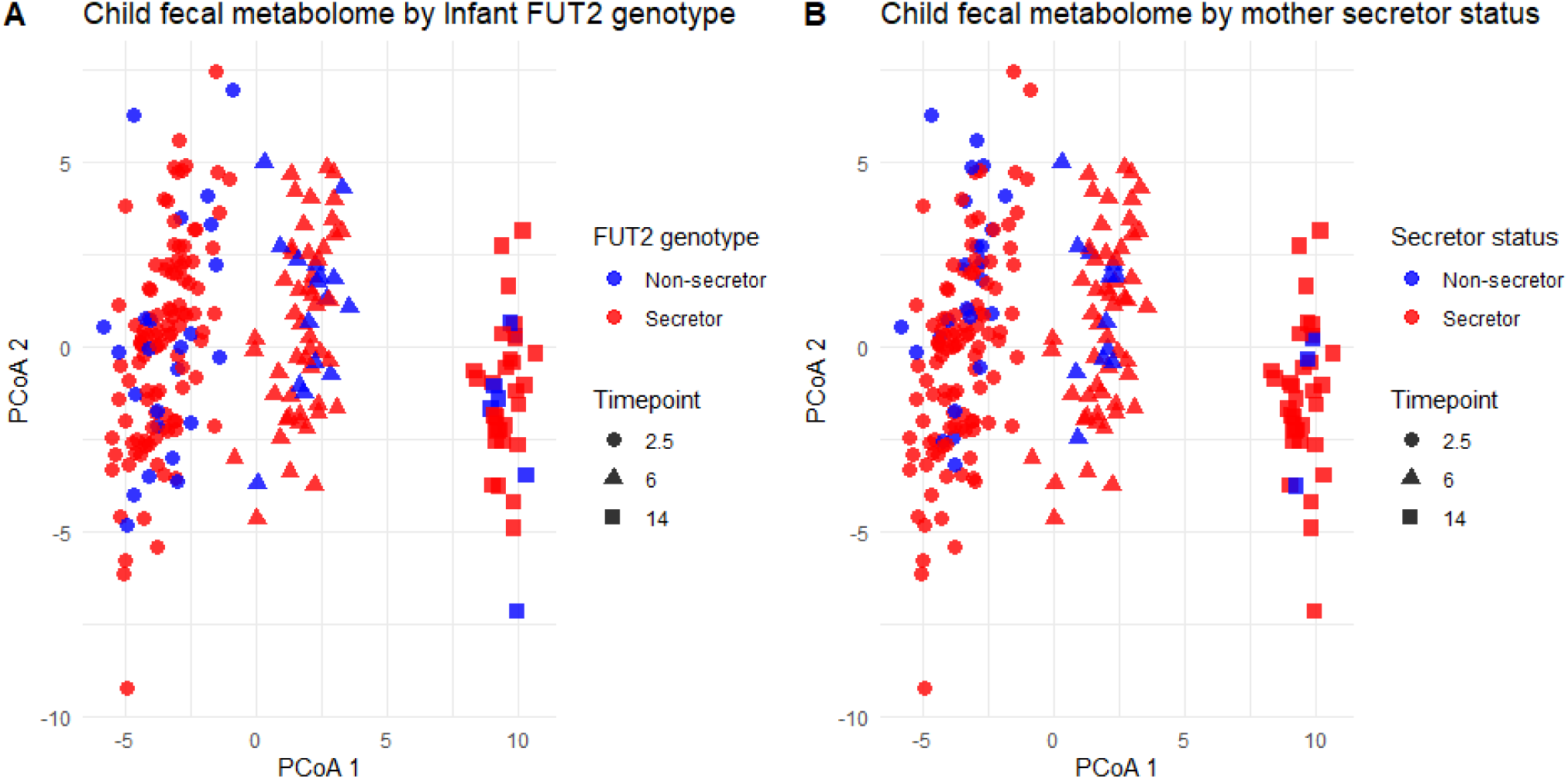
Dissimilarity of child fecal metabolome (Euclidean) in a principal component analysis by A) infant FUT2 genotype (secretor status) and B) phenotype of mother secretor status across timepoints.

**Figure S5.**
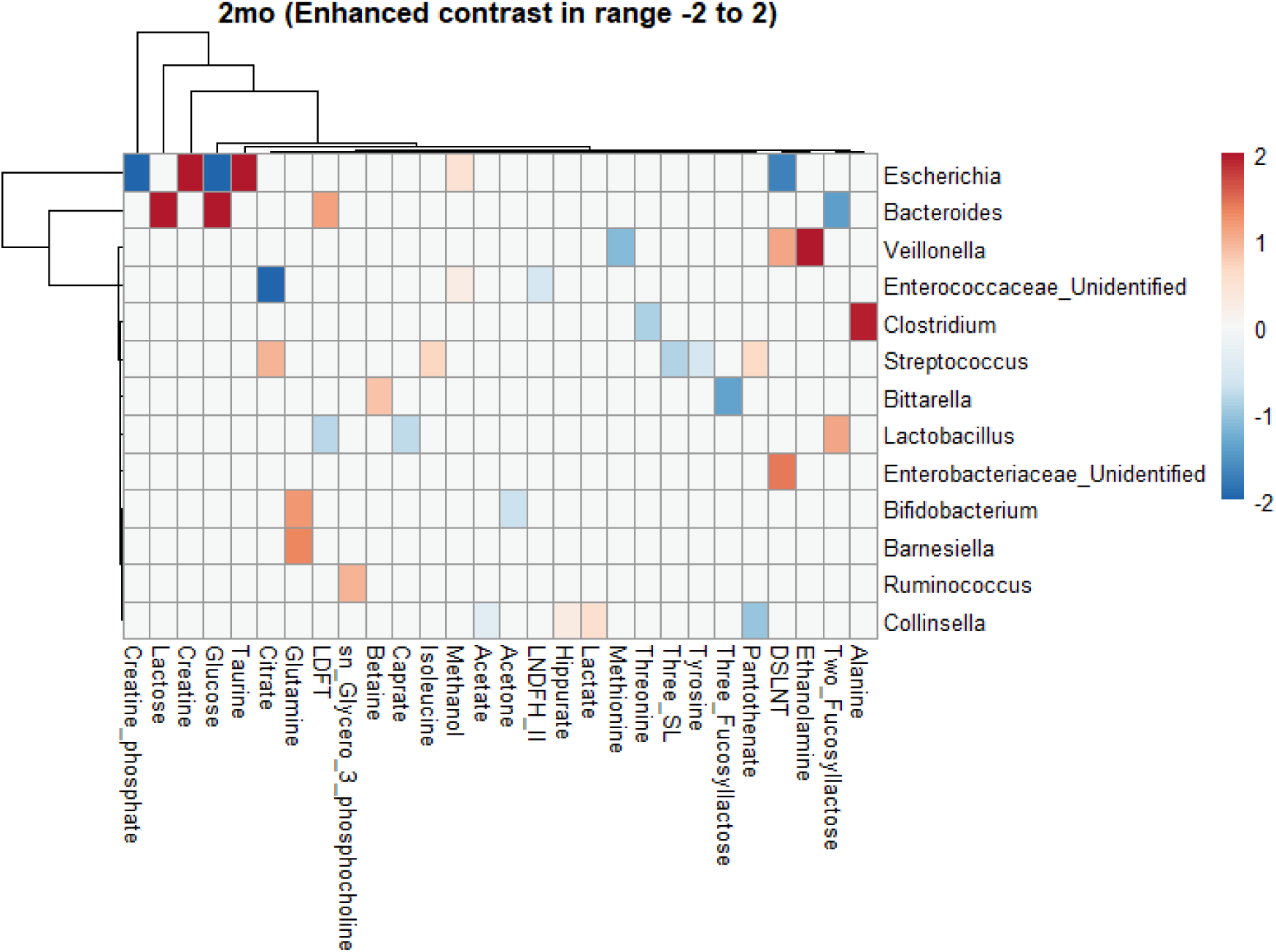
Differential abundance with milk metabolites in 2mo timepoint. Adjusted for covariates in linear model.

**Figure S6.**
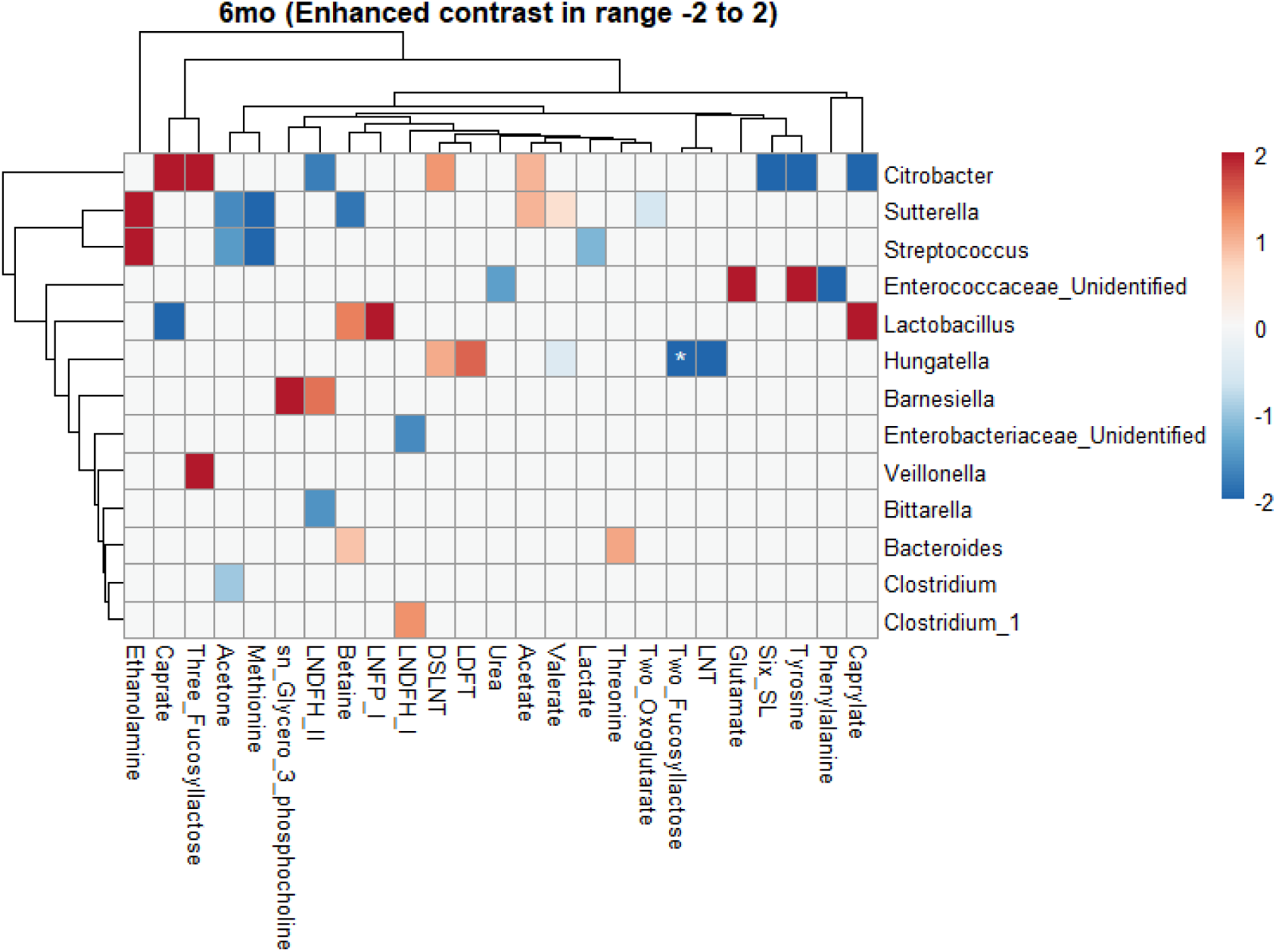
Differential abundance with milk metabolites in 6mo timepoint. Asterisk (*) adjusted p-value =0.09.

**Figure S7.**
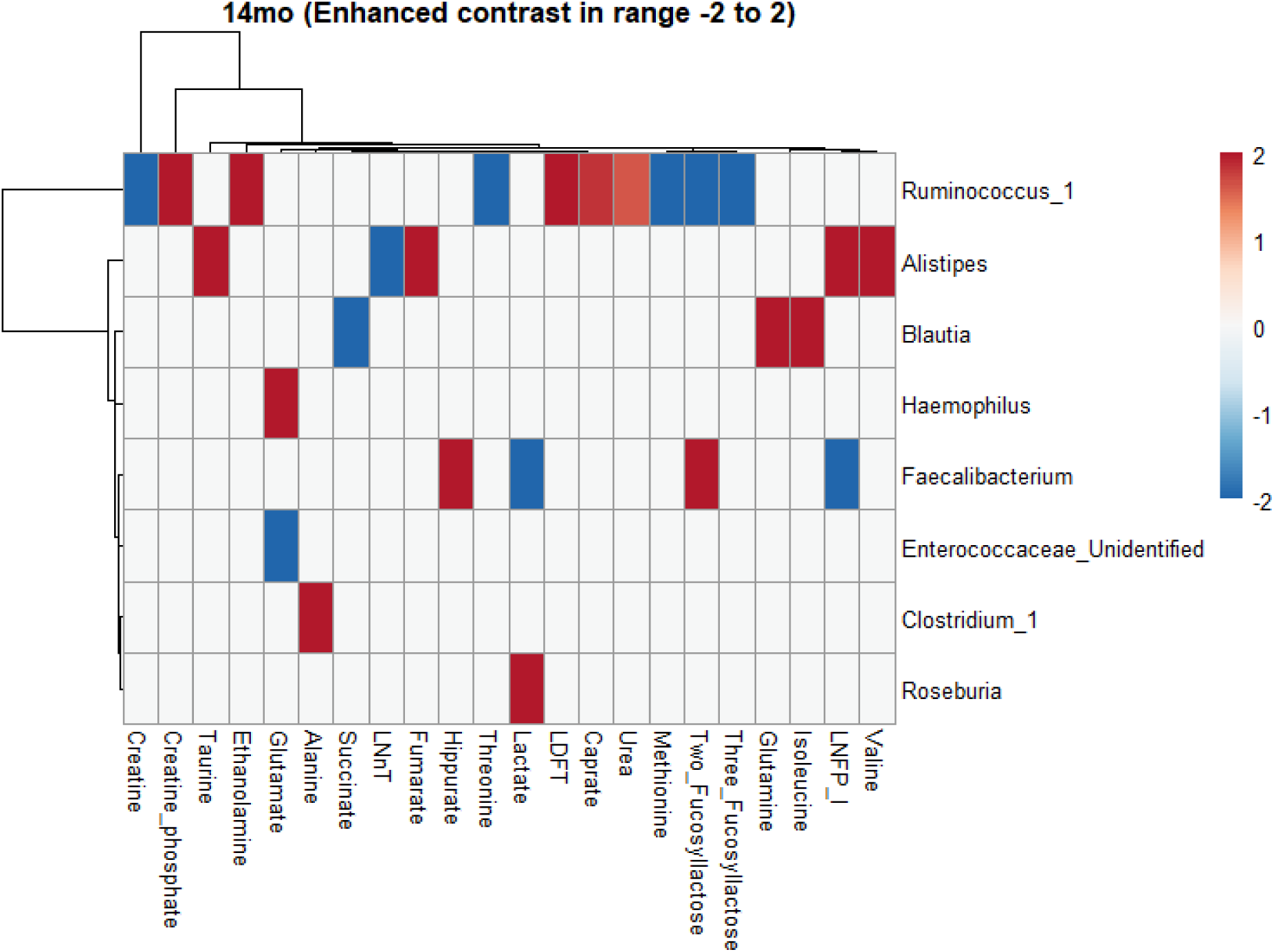
Differential abundance with milk metabolites in 14mo timepoint.

**Figure S8.**
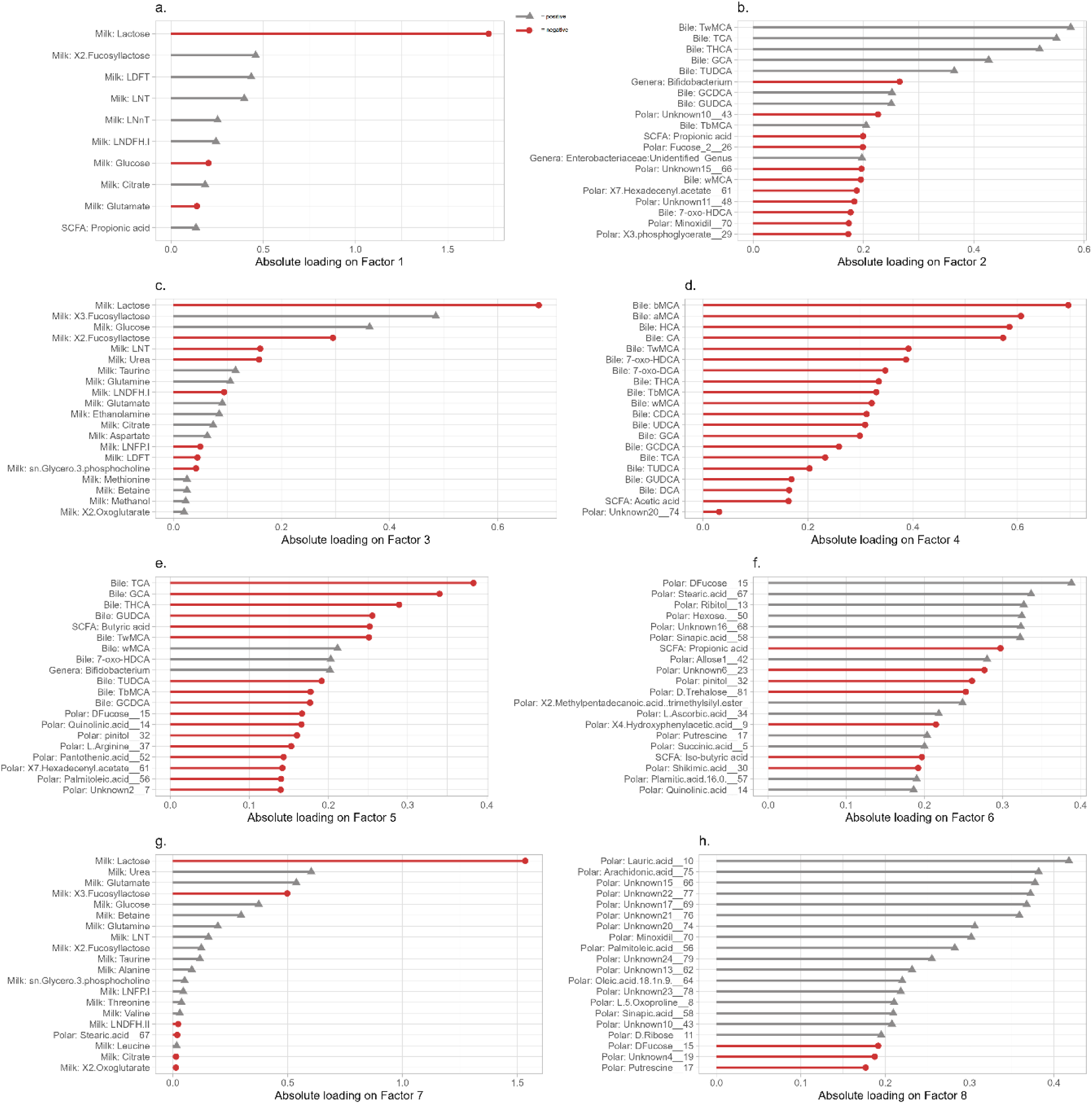
Factor loadings of MOFA. 8 factors with top20 loadings. Direction of loading indicated with colors and shapes (grey triangle= positive, red circle= negative).

**Figure S9.**
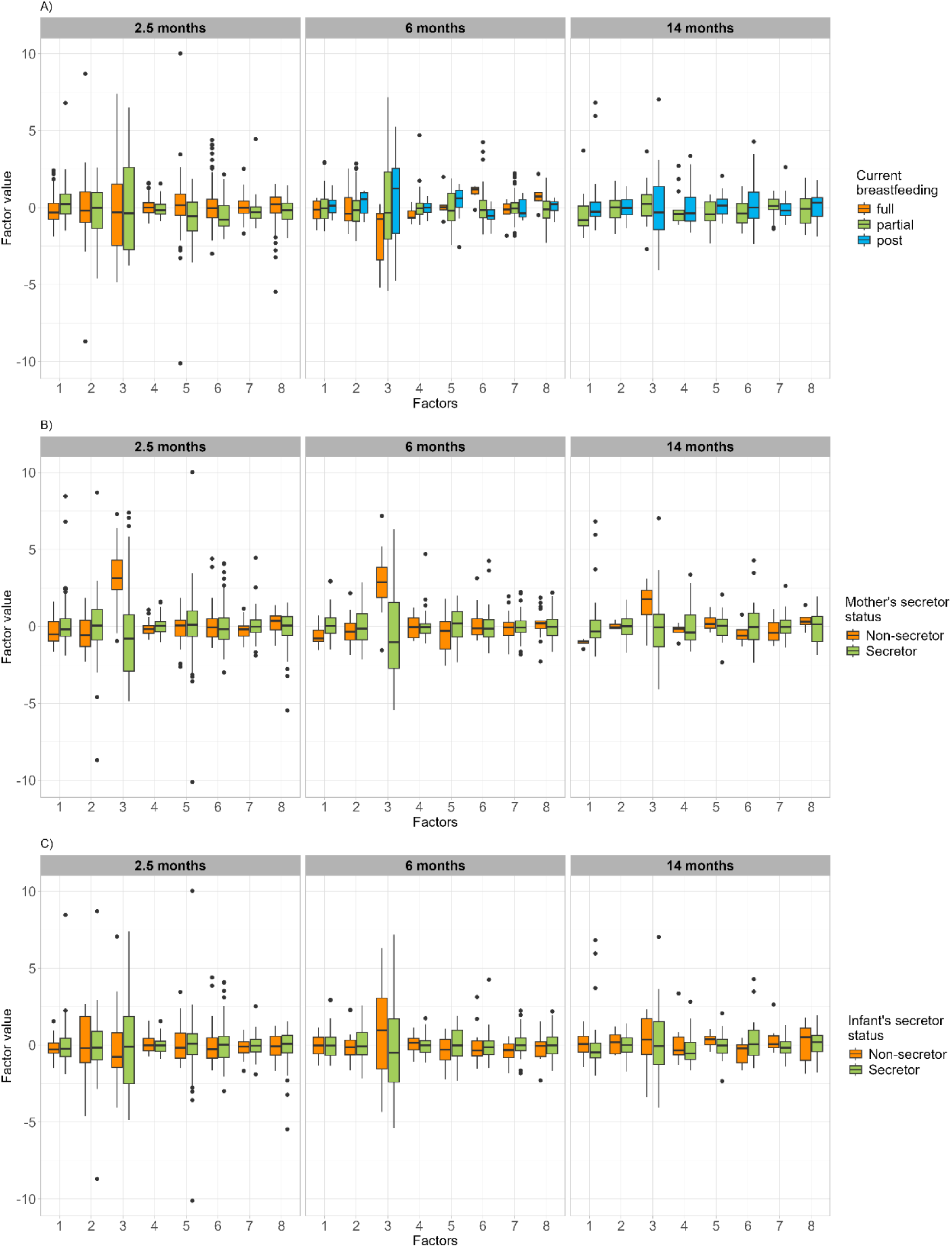
Factor values with breast feeding variables in three timepoints. Post in current feeding denotes recently ceased breastfeeding or the late stages of weaning. A) Boxplots of factor values by current feeding mode. Post indicates here that breastfeeding has recently ended. B) Boxplots of factor values by mother secretor status (Phenotype). C) Boxplots of factor values by infant secretor status (FUT2 genotype).

## References

1. Suárez-Martínez C, Santaella-Pascual M, Yagüe-Guirao G, Martínez-Graciá C. Infant gut microbiota colonization: influence of prenatal and postnatal factors, focusing on diet. Front Microbiol. 2023;14. 10.3389/fmicb.2023.1236254.

2. Dominguez-Bello MG, Godoy-Vitorino F, Knight R, Blaser MJ. Role of the microbiome in human development. Gut. 2019;68:1108–14. 10.1136/gutjnl-2018-317503.

3. Stewart CJ, Ajami NJ, O’Brien JL, Hutchinson DS, Smith DP, Wong MC, et al. Temporal development of the gut microbiome in early childhood from the TEDDY study. Nature. 2018;562:583–8. 10.1038/s41586-018-0617-x.

4. Azad MB, Konya T, Maughan H, Guttman DS, Field CJ, Chari RS, et al. Gut microbiota of healthy Canadian infants: profiles by mode of delivery and infant diet at 4 months. CMAJ. 2013;185:385–94. 10.1503/cmaj.121189.

5. Ames SR, Lotoski LC, Azad MB. Comparing early life nutritional sources and human milk feeding practices: personalized and dynamic nutrition supports infant gut microbiome development and immune system maturation. Gut Microbes. 2023;15:2190305. 10.1080/19490976.2023.2190305.

6. Korpela K, Salonen A, Hickman B, Kunz C, Sprenger N, Kukkonen K, et al. Fucosylated oligosaccharides in mother’s milk alleviate the effects of caesarean birth on infant gut microbiota. Sci Rep. 2018;8:13757. 10.1038/s41598-018-32037-6.

7. Kunz C, Meyer C, Collado MC, Geiger L, García-Mantrana I, Bertua-Ríos B, et al. Influence of Gestational Age, Secretor, and Lewis Blood Group Status on the Oligosaccharide Content of Human Milk. Journal of Pediatric Gastroenterology and Nutrition. 2017;64:789–98. 10.1097/MPG.0000000000001402.

8. Brink LR, Mercer KE, Piccolo BD, Chintapalli SV, Elolimy A, Bowlin AK, et al. Neonatal diet alters fecal microbiota and metabolome profiles at different ages in infants fed breast milk or formula. The American Journal of Clinical Nutrition. 2020;111:1190–202. 10.1093/ajcn/nqaa076.

9. Lordan C, Roche AK, Delsing D, Nauta A, Groeneveld A, MacSharry J, et al. Linking human milk oligosaccharide metabolism and early life gut microbiota: bifidobacteria and beyond. Microbiology and Molecular Biology Reviews. 2024;88:e00094–23. 10.1128/mmbr.00094-23.

10. Yelverton CA, Killeen SL, Feehily C, Moore RL, Callaghan SL, Geraghty AA, et al. Maternal breastfeeding is associated with offspring microbiome diversity; a secondary analysis of the MicrobeMom randomized control trial. Front Microbiol. 2023;14. 10.3389/fmicb.2023.1154114.

11. Porro M, Kundrotaite E, Mellor DD, Munialo CD. A narrative review of the functional components of human breast milk and their potential to modulate the gut microbiome, the consideration of maternal and child characteristics, and confounders of breastfeeding, and their impact on risk of obesity later in life. Nutrition Reviews. 2023;81:597–609. 10.1093/nutrit/nuac072.

12. Jokela R, Ponsero AJ, Dikareva E, Wei X, Kolho K-L, Korpela K, et al. Sources of gut microbiota variation in a large longitudinal Finnish infant cohort. eBioMedicine. 2023;94:104695. 10.1016/j.ebiom.2023.104695.

13. Hickman B, Salonen A, Ponsero AJ, Jokela R, Kolho K-L, de Vos WM, et al. Gut microbiota wellbeing index predicts overall health in a cohort of 1000 infants. Nat Commun. 2024;15:8323. 10.1038/s41467-024-52561-6.

14. Matharu D, Ponsero AJ, Lengyel M, Meszaros-Matwiejuk A, Kolho K-L, de Vos WM, et al. Human milk oligosaccharide composition is affected by season and parity and associates with infant gut microbiota in a birth mode dependent manner in a Finnish birth cohort. EBioMedicine. 2024;104:105182. 10.1016/j.ebiom.2024.105182.

15. Roager HM, Stanton C, Hall LJ. Microbial metabolites as modulators of the infant gut microbiome and host-microbial interactions in early life. Gut Microbes. 2023;15:2192151. 10.1080/19490976.2023.2192151.

16. Zhang Y, Chen R, Zhang D, Qi S, Liu Y. Metabolite interactions between host and microbiota during health and disease: Which feeds the other? Biomedicine & Pharmacotherapy. 2023;160:114295. 10.1016/j.biopha.2023.114295.

17. Liu J, Tan Y, Cheng H, Zhang D, Feng W, Peng C. Functions of Gut Microbiota Metabolites, Current Status and Future Perspectives. Aging Dis. 2022;13:1106–26. 10.14336/AD.2022.0104.

18. van Best N, Rolle-Kampczyk U, Schaap FG, Basic M, Olde Damink SWM, Bleich A, et al. Bile acids drive the newborn’s gut microbiota maturation. Nat Commun. 2020;11:3692. 10.1038/s41467-020-17183-8.

19. Hoen AG, Coker MO, Madan JC, Pathmasiri W, McRitchie S, Dade EF, et al. Association of Cesarean Delivery and Formula Supplementation with the Stool Metabolome of 6-Week-Old Infants. Metabolites. 2021;11:702. 10.3390/metabo11100702.

20. Holzhausen EA, Shen N, Chalifour B, Tran V, Li Z, Sarnat JA, et al. Longitudinal profiles of the fecal metabolome during the first 2 years of life. Sci Rep. 2023;13:1886. 10.1038/s41598-023-28862-z.

21. Bode L. Human milk oligosaccharides: every baby needs a sugar mama. Glycobiology. 2012;22:1147–62. 10.1093/glycob/cws074.

22. Kashyap PC, Marcobal A, Ursell LK, Smits SA, Sonnenburg ED, Costello EK, et al. Genetically dictated change in host mucus carbohydrate landscape exerts a diet-dependent effect on the gut microbiota. Proceedings of the National Academy of Sciences. 2013;110:17059–64. 10.1073/pnas.1306070110.

23. Wacklin P, Mäkivuokko H, Alakulppi N, Nikkilä J, Tenkanen H, Räbinä J, et al. Secretor Genotype (FUT2 gene) Is Strongly Associated with the Composition of Bifidobacteria in the Human Intestine. PLOS ONE. 2011;6:e20113. 10.1371/journal.pone.0020113.

24. Wacklin P, Tuimala J, Nikkilä J, Tims S, Mäkivuokko H, Alakulppi N, et al. Faecal Microbiota Composition in Adults Is Associated with the FUT2 Gene Determining the Secretor Status. PLOS ONE. 2014;9:e94863. 10.1371/journal.pone.0094863.

25. Karlsson L, Tolvanen M, Scheinin NM, Uusitupa H-M, Korja R, Ekholm E, et al. Cohort Profile: The FinnBrain Birth Cohort Study (FinnBrain). International Journal of Epidemiology. 2018;47:15–16j. 10.1093/ije/dyx173.

26. Cox JL, Holden JM, Sagovsky R. Detection of Postnatal Depression: Development of the 10-item Edinburgh Postnatal Depression Scale. The British Journal of Psychiatry. 1987;150:782–6. 10.1192/bjp.150.6.782.

27. Derogatis LR, Lipman RS, Covi L. SCL-90: an outpatient psychiatric rating scale--preliminary report. Psychopharmacol Bull. 1973;9:13–28.

28. Sundekilde UK, Downey E, O’Mahony JA, O’Shea C-A, Ryan CA, Kelly AL, et al. The Effect of Gestational and Lactational Age on the Human Milk Metabolome. Nutrients. 2016;8:304. 10.3390/nu8050304.

29. Dror DK, Allen LH. Overview of Nutrients in Human Milk. Advances in Nutrition. 2018;9:278S–294S. 10.1093/advances/nmy022.

30. Korhonen LS, Lukkarinen M, Kantojärvi K, Räty P, Karlsson H, Paunio T, et al. Interactions of genetic variants and prenatal stress in relation to the risk for recurrent respiratory infections in children. Sci Rep. 2021;11:7589. 10.1038/s41598-021-87211-0.

31. Acosta H, Kantojärvi K, Tuulari JJ, Lewis JD, Hashempour N, Scheinin NM, et al. Association of cumulative prenatal adversity with infant subcortical structure volumes and child problem behavior and its moderation by a coexpression polygenic risk score of the serotonin system. Development and Psychopathology. 2024;36:1027–42. 10.1017/S0954579423000275.

32. Loh P-R, Danecek P, Palamara PF, Fuchsberger C, A Reshef Y, K Finucane H, et al. Reference-based phasing using the Haplotype Reference Consortium panel. Nat Genet. 2016;48:1443–8. 10.1038/ng.3679.

33. Browning BL, Browning SR. Genotype Imputation with Millions of Reference Samples. The American Journal of Human Genetics. 2016;98:116–26. 10.1016/j.ajhg.2015.11.020.

34. Rintala A, Pietilä S, Munukka E, Eerola E, Pursiheimo J-P, Laiho A, et al. Gut Microbiota Analysis Results Are Highly Dependent on the 16S rRNA Gene Target Region, Whereas the Impact of DNA Extraction Is Minor. J Biomol Tech. 2017;28:19–30. 10.7171/jbt.17-2801-003.

35. Callahan BJ, McMurdie PJ, Rosen MJ, Han AW, Johnson AJA, Holmes SP. DADA2: High-resolution sample inference from Illumina amplicon data. Nat Methods. 2016;13:581–3. 10.1038/nmeth.3869.

36. Quast C, Pruesse E, Yilmaz P, Gerken J, Schweer T, Yarza P, et al. The SILVA ribosomal RNA gene database project: improved data processing and web-based tools. Nucleic Acids Res. 2013;41 Database issue:D590-596. 10.1093/nar/gks1219.

37. Yilmaz P, Parfrey LW, Yarza P, Gerken J, Pruesse E, Quast C, et al. The SILVA and “All-species Living Tree Project (LTP)” taxonomic frameworks. Nucleic Acids Research. 2014;42:D643–8. 10.1093/nar/gkt1209.

38. Wang Q, Garrity GM, Tiedje JM, Cole JR. Naïve Bayesian Classifier for Rapid Assignment of rRNA Sequences into the New Bacterial Taxonomy. Applied and Environmental Microbiology. 2007;73:5261–7. 10.1128/AEM.00062-07.

39. Aatsinki A-K, Lamichhane S, Isokääntä H, Sen P, Kråkström M, Alves MA, et al. Dynamics of Gut Metabolome and Microbiome Maturation during Early Life. 2023;:2023.05.29.23290441. 10.1101/2023.05.29.23290441.

40. Trimigno A, Khakimov B, Mejia JLC, Mikkelsen MS, Kristensen M, Jespersen BM, et al. Identification of weak and gender specific effects in a short 3 weeks intervention study using barley and oat mixed linkage β-glucan dietary supplements: a human fecal metabolome study by GC-MS. Metabolomics. 2017;13:108. 10.1007/s11306-017-1247-2.

41. Lamichhane S, Sen P, Dickens AM, Orešič M, Bertram HC. Gut metabolome meets microbiome: A methodological perspective to understand the relationship between host and microbe. Methods. 2018;149:3–12. 10.1016/j.ymeth.2018.04.029.

42. R: The R Project for Statistical Computing. https://www.r-project.org/. Accessed 14 Oct 2024.

43. Kassambara A, Mundt F. factoextra: Extract and Visualize the Results of Multivariate Data Analyses. cran.r-project.org/web/packages/factoextra/index.html. 2020.

44. Borman, Ernst, Lahti. https://github.com/microbiome/mia. 2024.

45. Oksanen J, Simpson GL, Blanchet FG, Kindt R, Legendre P, Minchin PR, et al. vegan: Community Ecology Package. 2001;:2.6–8. 10.32614/CRAN.package.vegan.

46. Davison AC, Hinkley DV. Bootstrap Methods and their Application. Cambridge: Cambridge University Press; 1997. 10.1017/CBO9780511802843.

47. Argelaguet R, Velten B, Arnol D, Dietrich S, Zenz T, Marioni JC, et al. Multi-Omics Factor Analysis-a framework for unsupervised integration of multi-omics data sets. Mol Syst Biol. 2018;14:e8124. 10.15252/msb.20178124.

48. Argelaguet R, Arnol D, Bredikhin D, Deloro Y, Velten B, Marioni JC, et al. MOFA+: a statistical framework for comprehensive integration of multi-modal single-cell data. Genome Biol. 2020;21:111. 10.1186/s13059-020-02015-1.

49. Velten B, Braunger JM, Argelaguet R, Arnol D, Wirbel J, Bredikhin D, et al. Identifying temporal and spatial patterns of variation from multimodal data using MEFISTO. Nat Methods. 2022;19:179–86. 10.1038/s41592-021-01343-9.

50. Wickham H. ggplot2. Cham: Springer International Publishing; 2016. 10.1007/978-3-319-24277-4.

51. Finnish food authority. Ruokavirasto. 2025. https://www.ruokavirasto.fi/elintarvikkeet/terveytta-edistava-ruokavalio/ravitsemus--ja-ruokasuositukset/imevaisikaiset-ja-lapset/. Accessed 11 Apr 2025.

52. Pelto J, Auranen K, Kujala JV, Lahti L. Elementary methods provide more replicable results in microbial differential abundance analysis. Brief Bioinform. 2025;26:bbaf130. 10.1093/bib/bbaf130.

53. Fernandes AD, Macklaim JM, Linn TG, Reid G, Gloor GB. ANOVA-Like Differential Expression (ALDEx) Analysis for Mixed Population RNA-Seq. PLOS ONE. 2013;8:e67019. 10.1371/journal.pone.0067019.

54. Soyyılmaz B, Mikš MH, Röhrig CH, Matwiejuk M, Meszaros-Matwiejuk A, Vigsnæs LK. The Mean of Milk: A Review of Human Milk Oligosaccharide Concentrations throughout Lactation. Nutrients. 2021;13:2737. 10.3390/nu13082737.

55. Azad MB, Wade KH, Timpson NJ. FUT2 secretor genotype and susceptibility to infections and chronic conditions in the ALSPAC cohort. Wellcome Open Res. 2018;3:65. 10.12688/wellcomeopenres.14636.2.

56. Sabater C, Iglesias-Gutiérrez E, Ruiz L, Margolles A. Next-generation sequencing of the athletic gut microbiota: a systematic review. Microbiome Res Rep. 2023;2:5. 10.20517/mrr.2022.16.

57. Nair MKM, Joy J, Vasudevan P, Hinckley L, Hoagland TA, Venkitanarayanan KS. Antibacterial Effect of Caprylic Acid and Monocaprylin on Major Bacterial Mastitis Pathogens. Journal of Dairy Science. 2005;88:3488–95. 10.3168/jds.S0022-0302(05)73033-2.

58. Han Q, Liu R, Wang H, Zhang R, Liu H, Li J, et al. Gut Microbiota-Derived 5-Hydroxyindoleacetic Acid Alleviates Diarrhea in Piglets via the Aryl Hydrocarbon Receptor Pathway. J Agric Food Chem. 2023;71:15132–44. 10.1021/acs.jafc.3c04658.

59. Li Z, Zhu Y, Ni D, Zhang W, Mu W. Occurrence, functional properties, and preparation of 3-fucosyllactose, one of the smallest human milk oligosaccharides. Crit Rev Food Sci Nutr. 2023;63:9364–78. 10.1080/10408398.2022.2064813.

60. Hill DR, Buck RH. Infants Fed Breastmilk or 2ʹ-FL Supplemented Formula Have Similar Systemic Levels of Microbiota-Derived Secondary Bile Acids. Nutrients. 2023;15:2339. 10.3390/nu15102339.

61. Gazi MA, Fahim SM, Hasan MM, Hossaini F, Alam MA, Hossain MS, et al. Maternal and child FUT2 and FUT3 status demonstrate relationship with gut health, body composition and growth of children in Bangladesh. Sci Rep. 2022;12:18764. 10.1038/s41598-022-23616-9.

62. Lewis ZT, Totten SM, Smilowitz JT, Popovic M, Parker E, Lemay DG, et al. Maternal fucosyltransferase 2 status affects the gut bifidobacterial communities of breastfed infants. Microbiome. 2015;3:13. 10.1186/s40168-015-0071-z.

63. Laursen MF. Gut Microbiota Development: Influence of Diet from Infancy to Toddlerhood. Ann Nutr Metab. 2021;:1–14. 10.1159/000517912.

64. Thorman AW, Adkins G, Conrey SC, Burrell AR, Yu Y, White B, et al. Gut Microbiome Composition and Metabolic Capacity Differ by FUT2 Secretor Status in Exclusively Breastfed Infants. Nutrients. 2023;15:471. 10.3390/nu15020471.

65. Wang A, Diana A, Rahmannia S, Gibson RS, Houghton LA, Slupsky CM. Impact of milk secretor status on the fecal metabolome and microbiota of breastfed infants. Gut Microbes. 2023;15:2257273. 10.1080/19490976.2023.2257273.

66. Sugino KY, Ma T, Kerver JM, Paneth N, Comstock SS. Human Milk Feeding Patterns at 6 Months of Age are a Major Determinant of Fecal Bacterial Diversity in Infants. J Hum Lact. 2021;37:703–13. 10.1177/0890334420957571.

67. Gurung M, Schlegel BT, Rajasundaram D, Fox R, Bode L, Yao T, et al. Microbiota from human infants consuming secretors or non-secretors mothers’ milk impacts the gut and immune system in mice. mSystems. 2024;9:e00294–24. 10.1128/msystems.00294-24.

68. Gao X, Wu D, Wen Y, Gao L, Liu D, Zhong R, et al. Antiviral effects of human milk oligosaccharides: A review. International Dairy Journal. 2020;110:104784. 10.1016/j.idairyj.2020.104784.

69. Jantscher-Krenn E, Zherebtsov M, Nissan C, Goth K, Guner YS, Naidu N, et al. The human milk oligosaccharide disialyllacto-N-tetraose prevents necrotising enterocolitis in neonatal rats. Gut. 2012;61:1417–25. 10.1136/gutjnl-2011-301404.

70. Jabeen I, Islam S, Hassan AKMI, Tasnim Z, Shuvo SR. A brief insight into Citrobacter species - a growing threat to public health. Front Antibiot. 2023;2. 10.3389/frabi.2023.1276982.

71. Spigaglia P. Clostridioides difficile and Gut Microbiota: From Colonization to Infection and Treatment. Pathogens. 2024;13:646. 10.3390/pathogens13080646.

72. Zhai L, Huang C, Ning Z, Zhang Y, Zhuang M, Yang W, et al. *Ruminococcus gnavus* plays a pathogenic role in diarrhea-predominant irritable bowel syndrome by increasing serotonin biosynthesis. Cell Host & Microbe. 2023;31:33–44.e5. 10.1016/j.chom.2022.11.006.

73. Thomson P, Medina DA, Garrido D. Human milk oligosaccharides and infant gut bifidobacteria: Molecular strategies for their utilization. Food Microbiology. 2018;75:37–46. 10.1016/j.fm.2017.09.001.

74. Chapman JA, Masi AC, Beck LC, Watson H, Young GR, Browne HP, et al. Human milk oligosaccharide metabolism by Clostridium species suppresses inflammation and pathogen growth. 2025;:2025.01.21.633585. 10.1101/2025.01.21.633585.

75. Charbonneau MR, O’Donnell D, Blanton LV, Totten SM, Davis JCC, Barratt MJ, et al. Sialylated Milk Oligosaccharides Promote Microbiota-Dependent Growth in Models of Infant Undernutrition. Cell. 2016;164:859–71. 10.1016/j.cell.2016.01.024.

76. Lewis ED, Richard C, Goruk S, Wadge E, Curtis JM, Jacobs RL, et al. Feeding a Mixture of Choline Forms during Lactation Improves Offspring Growth and Maternal Lymphocyte Response to Ex Vivo Immune Challenges. Nutrients. 2017;9:713. 10.3390/nu9070713.

77. Romano KA, Vivas EI, Amador-Noguez D, Rey FE. Intestinal Microbiota Composition Modulates Choline Bioavailability from Diet and Accumulation of the Proatherogenic Metabolite Trimethylamine-N-Oxide. mBio. 2015;6: 10.1128/mbio.02481-14. 10.1128/mbio.02481-14.

78. Seki D, Errerd T, Hall LJ. The role of human milk fats in shaping neonatal development and the early life gut microbiota. Microbiome Res Rep. 2023;2:8. 10.20517/mrr.2023.09.

79. Macfarlane GT, Macfarlane S. Bacteria, Colonic Fermentation, and Gastrointestinal Health. Journal of AOAC INTERNATIONAL. 2012;95:50–60. 10.5740/jaoacint.SGE_Macfarlane.

80. Rios-Covian D, González S, Nogacka AM, Arboleya S, Salazar N, Gueimonde M, et al. An Overview on Fecal Branched Short-Chain Fatty Acids Along Human Life and as Related With Body Mass Index: Associated Dietary and Anthropometric Factors. Front Microbiol. 2020;11. 10.3389/fmicb.2020.00973.

81. Yang S, Yu X, Zuo Q. Branched-Chain Fatty Acids and Obesity: A Narrative Review. Nutr Rev. 2025;83:1314–26. 10.1093/nutrit/nuaf022.

82. You X, Rani A, Özcan E, Lyu Y, Sela DA. Bifidobacterium longum subsp. infantis utilizes human milk urea to recycle nitrogen within the infant gut microbiome. Gut Microbes. 2023;15:2192546. 10.1080/19490976.2023.2192546.

83. Firth IJ, Sim MAR, Fitzgerald BG, Moore AE, Pittao CR, Gianetto-Hill C, et al. Urease in acetogenic Lachnospiraceae drives urea carbon salvage in SCFA pools. Gut Microbes. 2025;17:2492376. 10.1080/19490976.2025.2492376.

84. Ribo S, Sánchez-Infantes D, Martinez-Guino L, García-Mantrana I, Ramon-Krauel M, Tondo M, et al. Increasing breast milk betaine modulates Akkermansia abundance in mammalian neonates and improves long-term metabolic health. Sci Transl Med. 2021;13:eabb0322. 10.1126/scitranslmed.abb0322.

85. Li Z, Hu G, Zhu L, Sun Z, Jiang Y, Gao M, et al. Study of growth, metabolism, and morphology of Akkermansia muciniphila with an in vitro advanced bionic intestinal reactor. BMC Microbiology. 2021;21:61. 10.1186/s12866-021-02111-7.

86. Poulsen KO, Meng F, Lanfranchi E, Young JF, Stanton C, Ryan CA, et al. Dynamic Changes in the Human Milk Metabolome Over 25 Weeks of Lactation. Front Nutr. 2022;9. 10.3389/fnut.2022.917659.

87. Schoeler M, Caesar R. Dietary lipids, gut microbiota and lipid metabolism. Rev Endocr Metab Disord. 2019;20:461–72. 10.1007/s11154-019-09512-0.

88. Yokota A, Fukiya S, Islam KBMS, Ooka T, Ogura Y, Hayashi T, et al. Is bile acid a determinant of the gut microbiota on a high-fat diet? Gut Microbes. 2012;3:455–9. 10.4161/gmic.21216.

89. Tessier MEM, Shneider BL, Petrosino JF, Preidis GA. Bile acid and microbiome interactions in the developing child. J Pediatr Gastroenterol Nutr. 2025;80:832–9. 10.1002/jpn3.70014.

90. Blaak EE, Canfora EE, Theis S, Frost G, Groen AK, Mithieux G, et al. Short chain fatty acids in human gut and metabolic health. 2020. 10.3920/BM2020.0057.

91. Hsu C-Y, Khachatryan LG, Younis NK, Mustafa MA, Ahmad N, Athab ZH, et al. Microbiota-derived short chain fatty acids in pediatric health and diseases: from gut development to neuroprotection. Front Microbiol. 2024;15. 10.3389/fmicb.2024.1456793.

